# A cryopreservation strategy for myoblast storage in paper-based scaffolds for inter-laboratory studies of skeletal muscle health

**DOI:** 10.1101/2024.03.07.583636

**Authors:** Saifedine T. Rjaibi, Erik Jacques, Jiaru Ni, Bin Xu, Sonya Kouthouridis, Julie Sitolle, Heta Lad, Nitya Gulati, Nancy T. Li, Henry Ahn, Howard J. Ginsberg, Boyang Zhang, Fabien Le Grand, Penney M. Gilbert, Alison P. McGuigan

## Abstract

Three-dimensional tissue-engineered models are poised to facilitate understanding of skeletal muscle pathophysiology and identify novel therapeutic agents to improve muscle health. Adopting these culture models within the broader biology community is a challenge as many models involve complex methodologies and significant investments of time and resources to optimize manufacturing protocols. To alleviate this barrier, we developed a protocol with commercially available reagents to cryopreserve myoblasts in a 96-well compatible format that allows tissues to be transferred to users without expertise in 2D or 3D skeletal muscle cell culture. We validate that myoblasts encapsulated in a hydrogel and cryopreserved in paper-based scaffolds maintain cell viability, differentiation, and function via acetylcholine-induced transient calcium responses. Furthermore, we demonstrate successful shipping of myoblasts cryopreserved in paper-based scaffolds to intra-provincial and international collaborators who successfully thawed, cultured, and used the 3D muscle tissues. Finally, we confirm the application of our method to study muscle endogenous repair by seeding freshly isolated skeletal muscle stem cells to cryopreserved then differentiated and injured tissues, demonstrating expected responses to a known stimulator of muscle stem cell self-renewal, p38α/β MAPKi. Altogether, our 3D myoblast cryopreservation protocol offers broadened access of a complex skeletal muscle tissue model to the research community.

## Introduction

Skeletal muscle powers everyday functions of the human body from locomotion, metabolism, and thermogenesis.[1] However, numerous conditions can disrupt normal skeletal muscle homeostasis, ranging from muscular dystrophies, aging, obesity, and pharmacological toxicity.[2–5] For many of these diseases, underlying pathological mechanisms continue to be poorly understood, and as a result, gold-standard treatments have remained elusive.

Animal models provide an important tool to understand skeletal muscle disease pathology and drug responses *in vivo,* in particular the intramuscular transplantation assay.[6–7] Advantages of animal models include their capacity for long-term studies and ability to model complex skeletal muscle biology due to the presence of the many cell types and the complex spatiotemporal gradients present in native muscle tissues. However, animal studies are often expensive, time-consuming, and require a high level of expertise. Further, interspecies differences between mice and humans combined with the low-throughput possible with such animal models make them an inefficient tool in the process of drug discovery.[8–9]

In vitro culture models derived from human myoblasts offer an alternative higher-throughput tool for studying human skeletal muscle biology. For example, two-dimensional (2D) models, such as myoblast monolayers on tissue culture plates, provide a simple platform that is relatively inexpensive, easy-to-use, and amenable to high-throughput drug screens. However, myotubes in 2D cultures lack alignment and the intrinsic three-dimensional architecture of in vivo tissues, making them infeasible for evaluating functional metrics such as contractile force or fatigue resistance. Furthermore, the stiff surfaces of standard plastic tissue culture plates cannot support concentric muscle contractions, which eventually leads to myotube detachment.[10–12] These constraints limit the skeletal muscle biology that can be explored using 2D culture platforms.[12–15]

To expand the biology that can be studied in vitro, a variety of three-dimensional (3D) skeletal muscle models have been developed.[16–23] Importantly, 3D models are engineered to provide the structural integrity necessary to support longer term myotube culture and to permit quantification of phenotypic metrics such as myotube morphology, contractile force, and calcium release kinetics. While lower throughput than 2D cultures, these models still offer an order of magnitude higher throughput relative to animal models while allowing researchers precise control over the culture conditions for probing specific molecules, mechanisms, and dynamics involved in muscle repair.[24–26] Our group previously reported a 3D culture model for studying Muscle Endogenous Repair (MEndR) that enabled the prediction of skeletal muscle endogenous repair modulators in a 24-well platform.[27] The MEndR assay involves the formation of a 3D myotube template followed by engraftment of muscle stem cells into the template. This skeletal muscle/stem cells co-culture tissue is then chemically injured, and the capacity of the muscle stem cells to mediate repair of the template is assessed by analyzing phenotypic readouts. Manufacturing of the MEndR assay myotube template involves re-suspending primary myoblasts in a fibrin/reconstituted basement membrane ECM, infiltrating this suspension into a thin cellulose scaffold and then culturing for 7 days under conditions that favour myoblast differentiation. This produces a thin, 3D sheet of multinucleated myotubes that we refer to as the “myotube template”. Freshly isolated muscle stem cells can then be engrafted within this template simply by pipetting them in solution onto the template surface and culturing for 24-48h to enable stem cell engraftment throughout the myotube template. This provides an engineered muscle tissue for the study of endogenous repair. Our previous work demonstrated pheno-copying of in vivo outcomes following treatment with small molecules such as p38α/β mitogen-activated protein kinase inhibitor (MAPKi) and epidermal growth factor receptor inhibitor (EGFRi), both known modulators of muscle stem cell-mediated repair.[7, 28–31]

Beyond endogenous repair, MEndR is a potentially useful platform for studying a variety of questions related to skeletal muscle biology. Specifically, the geometry of the platform facilitates live or end-point imaging and characterization of both the myotubes and engrafted stem cell populations, which is challenging in many of the other typically thicker 3D models available. This makes the MEndR myotube template highly useful for probing questions related to muscle regeneration, toxicity screens, or template self-repair. While our initial model was low throughput, our group recently miniaturized MEndR into a 96-well footprint (“mini-MEndR”),[32] making it a potentially useful secondary screening tool for stratifying a greater number of “hits” from 2D high-throughput screens. Still, the capacity to robustly manufacture myotube templates has remained a major barrier limiting the broader adoption of this model. For example, we previously noted that 3-5 seeding attempts were required to master the manufacturing of mini-MEndR myotube templates by new users already proficient in myoblast 2D cell culture.[32] Therefore, we set out to establish a protocol to cryopreserve myotube templates after manufacturing to enable their distribution to the research community with a view to improving platform adoption. Specifically, we built upon previous work from Bashir and colleagues, who demonstrated that myoblasts encapsulated in a hydrogel mixture seeded onto 3D-printed polyethylene glycol dimethacrylate (PEGDMA) molds could be cryopreserved and successfully maintain structure and function upon thawing.[33] By adapting their approach, here we develop a standard operating procedure (SOP) for the cryopreservation of myoblasts in paper-based scaffolds. We show that myoblasts encapsulated in a fibrin/reconstituted basement membrane and cryopreserved in cellulose scaffolds maintain cell viability, differentiation, and function via a transient calcium response to acetylcholine stimulus. Further, we demonstrate that muscle stem cells engraft into myotube templates derived from frozen myoblasts and that treatment with a known stimulator of muscle stem cell-mediated skeletal muscle repair recapitulates the expected effects on the engineered tissue. Finally, we demonstrate proof-of-concept that cryopreserved myotube template pre-cursors can be shipped, thawed, cultured, and analyzed by new users in both domestic and international locations. We anticipate our cryopreservation protocol will facilitate broader adoption of the mini-MEndR platform to the research community by offering an easy-to-learn, off-the-shelf model for inter-laboratory studies of skeletal muscle health.

## Materials and Methods

### Ethics

Exactly as detailed in,[34] the collection and use of human skeletal muscle tissue was reviewed and approved by the Providence St. Joseph’s and St. Michael’s Healthcare Research Ethics Board (REB# 13-370) and the University of Toronto Office of Research Ethics reviewed the approved study and further assigned administrative approval (Protocol# 30754). All procedures in this study were performed in accordance with the guidelines and regulations of the respective Research Ethics Boards. Written consent was obtained from donors prior to the scheduled surgical procedure. Human skeletal muscle tissues removed from the multifidus muscle of patients undergoing lumbar spine surgery and designated for disposal were utilized in this study. Recruitment began August 18, 2014 and ended Aug 18, 2020. All collection and use of mouse skeletal muscle tissue in this study was conducted as described in the approved animal use protocols (#20012838), which was reviewed and approved by the local Animal Care Committee (ACC) within the Division of Comparative Medicine (DCM) at the University of Toronto. More broadly, all animal use was conducted in accordance with the guidelines and regulations of the Canadian Council on Animal Care.

### Primary human myoblast maintenance

The STEM21 human primary myoblast culture was established as previously described.[22] For primary myoblast maintenance, plastic tissue culture dishes (Sarstedt, 100 x 20mm, 83.3902) were coated with collagen (Gibco, #A10483-01) and stored at 4 °C. Immediately before thawing cells, the collagen solution was aspirated, and the plate was washed once with PBS and allowed to dry in the biosafety cabinet. Next, a cryovial of primary human myoblasts (STEM21, passage 7-8) was quickly thawed in a 37 °C water bath and the contents of the vial diluted 10X in wash media (Table 1). The diluted cells were centrifuged, the supernatant aspirated, and the cell pellet re-suspended in wash media. 8 mL growth media (GM; Table 1) was dispensed into each 10 cm plate, to which the cells were then distributed evenly at a density of 5000-7000 cells/cm^2^ (manufacturer’s recommendation). Culture dishes were maintained in a cell culture incubator at 37 °C and 5 % CO2. Cells were maintained in GM until reaching 70-80 % confluency. Culture media was replaced every two days.

**Table 1.** Culture media formulations and solutions.

|  |  |
| --- | --- |
| Wash Medium | ● 89% DMEM (Gibco, #11995-065)<br>● 10% Fetal bovine serum (FBS) (Gibco, 12483-020)<br>● 1% Penicillin-Streptomycin (Gibco, #15140-122) |
| 2D Primary Human Myoblast Growth Medium (GM) | ● Ham's F-10 nutrient mix (Wisent 318-050-CL)<br>● 20% Fetal bovine serum (FBS)<br>● 1% Pen-strep (P/S)<br>● 5 ng/mL rh-FGF2 (ImmunoTools, 11343625) |
| 2D Primary Mouse Myoblast Growth Medium (SAT-10) | ● DMEM/F12 (Gibco, #11320-033)<br>● 1% Pen-strep (P/S)<br>● 20% Fetal bovine serum (FBS)<br>● 10% Horse serum (Gibco, #16050-122)<br>● 1% Glutamax (Gibco, #35050-061)<br>● 1% Insulin-transferrin-selenium (Gibco, #41400-045)<br>● 1% Non-essential amino acids (Gibco, #11140-050)<br>● 1% Sodium pyruvate (Gibco, #11360-070)<br>● 50 μM β-mercaptoethanol (Gibco, #21985-023)<br>● 5 ng/mL rh-FGF2 |
| Primary Human Myoblast Seeding Medium (SM) | ● GM without added FGF2<br>● 1.5 mg/mL Aminocaproic acid (ACA) (Sigma, A2504) |
| Primary Mouse Myoblast Seeding Medium (SAT-10 (-)FGF) | ● SAT-10 without added FGF2<br>● 1.5 mg/mL Aminocaproic acid (ACA) |
| Differentiation Medium (DM) | ● DMEM<br>● 1% Pen-strep (P/S)<br>● 2% Horse serum<br>● 10 μg/mL Insulin (Sigma, #I6634)<br>● 2 mg/mL Aminocaproic acid (ACA) |
| Myotube Template Injury Medium | ● Differentiation Medium<br>● 0.5 μM Cardiotoxin |
| MEndR Assay Regeneration Medium | ● Differentiation Medium<br>● 5 ng/mL rh-FGF2<br>● 10 μM p38α/$\beta$ MAPKi (Drug Name: SB203580) (New England biolabs, #5633) |
| Freezing Medium | ● 90% FBS<br>● 10% Dimethyl sulfoxide (DMSO) (Sigma, D8418) |
| Fibrinogen Solution | ● 10 mg/mL Fibrinogen (Sigma, #F8630) in NaCl solution (0.9 % wt/vol in ddH<sub>2</sub>O) |
| Extracellular Matrix (ECM) Master Mix | ● 40% DMEM<br>● 40% Fibrinogen<br>● 20% Geltrex™ (Thermofisher, A1413202) |
| Blocking Solution | ● Phosphate-buffered saline (PBS)<br>● 10% goat serum (Gibco, #16210072)<br>● 0.3% Triton X-100 (BioShop, #TRX777) |

### Primary human induced pluripotent stem cell (iPSC)-derived myogenic progenitor maintenance

Human iPSC-derived myogenic progenitors cultures were obtained using the protocol published by Xi et al.[35] with minor modifications. Specifically, the iPSC-derived myogenic progenitors were maintained in SK Max medium (Wisent Bioproducts, 301-061-CL) supplemented with 10% FBS, 20 ng/mL FGF2, 0.5% P/S as we previously described.[36–37] Cells were seeded at about 5x10^4^ cells/cm^2^ onto GelTrex^TM^ coated tissue culture plates for 3-4 days prior to use in experiments.

### Primary mouse myoblast maintenance

Primary mouse myoblasts were derived and maintained as described in our previous work.[38] Briefly, primary mouse myoblasts were derived from enzymatically digested skeletal muscle tissue. After filtering and red blood cell lysis, the remaining mononucleated cell population was enriched via MACS for myogenic cells using the satellite cell isolation kit and subsequent integrin α-7 selection. The cells were then cultured on collagen-coated tissue culture plates in SAT-10 media (Table 1). Cells were passaged at 70% confluency and used from passage 5–9 for experiments.

### Paper scaffold candidate selection

Four paper scaffold candidates were selected from a larger pool of candidate cellulose-based papers based on thickness and similarity of the material to the original paper scaffold used in for the MEndR[27] and mini-MEndR[32] protocols (Mini-Minit Products, R10, Scarborough, Canada). The four candidates were (1) Finum (Tea Filters XL, Riensch & Held GmbH & Co.KG, Germany), (2) Teeli (Teeli Bag Filter Large Square, Riensch & Held GmbH & Co.KG, Germany), (3) Twin Rivers (TR Coffee FIL WS 17.0 CR NAT, Twin Rivers Paper Company, USA), and (4) Ahlstrom-Munksjo (Qualitative Filter Paper Grade 601, Ahlstrom-Munksjo Filtration LLC, Mount Holly Springs, PA, USA).

Each of the four paper candidates were characterized using scanning electron microscopy (SEM) to assess their microstructural properties and for their ability to support myoblast differentiation, and compared to the original paper scaffold.

### Scanning electron microscopy (SEM)

SEM images were obtained using a Hitachi SEM SU3500 (Hitachi High-Technologies Canada Inc., Toronto, Canada). Samples were mounted onto carbon-tape coated stubs and gold–palladium sputter coated for 55 s using a Bal-Tec SCD050 Sample Sputter Coater (Leica Biosystems, USA) and then imaged at 10 kVD.

To characterize the microstructure of candidate papers, SEM images were manually thresholded in FIJI to distinguish between the pores and paper fibers. A list of all pores and their areas was generated using FIJI’s Analyze Particles and aggregated into a histogram of the number of pores within a certain size range, the overall porosity (% area of the image occupied by the pores), and the coefficient of variation (CV) of the area occupied by pores of a certain size using code developed in Python 3.

### Mini-myotube template fabrication and culture

For the cellulose paper selection experiments, circular paper scaffolds with a diameter of 5 mm (20 mm^2^) were cut out using a biopsy punch (Miltex 5 mm biopsy punch, Integra, 400-4450910). For all other experiments, rectangular paper scaffolds of dimensions 5mm x 4mm (20 mm^2^) were cut out from tea bag papers using scissors. Once cut, paper scaffolds were autoclaved under a vacuum cycle to ensure sterility (sterilization time = 20 minutes, dry time = 10 minutes). 96-well footprint myotube templates were generated according to the previously described method.[32, 38] Briefly, 48-well or 96-well tissue culture plates (Sarstedt, 83.3924) were coated with 200 μL or 100 μL pluronic acid, respectively, and stored at 4 °C for at least one hour. The pluronic acid was aspirated and the plate allowed to dry in the biosafety cabinet. Paper scaffolds were pre-adsorbed with 4-5 μL of 0.8 U/mL thrombin (Sigma-Aldrich, #T6884) in the 96/48-well plate, and once again allowed to dry in the biosafety cabinet with the plate lid removed. In the meantime, myoblasts were trypsinized from their 2D culture dishes, centrifuged, and re-suspended in the Extracellular Matrix (ECM) Master Mix (Table 1). Once the paper scaffolds were dry, the cell-ECM solution was pipetted into the scaffolds at a concentration of 10^5^ cells (human) or 2.5x10^4^ cells (mouse) per 4-5 μL ECM Master Mix (Table 2). Both the plate and ECM solution were kept on ice during the seeding process to delay ECM hydrogel gelation. Once seeding was completed, the well plate was transferred to the cell culture incubator (37 °C, 5 % CO2) for 5-10 minutes to initiate hydrogel gelation. Next, 500 μL or 200 μL of Seeding Media (SM; Table 1) was added to each well of the 48-well or 96-well plate, respectively, and the plate was then returned to the incubator. For standard (unfrozen) conditions, tissues were incubated for 2 days in SM, at which point the media was switched to Differentiation Media (DM; Table 1) and half- media changes with fresh DM were performed every other day thereafter.

**Table 2.**
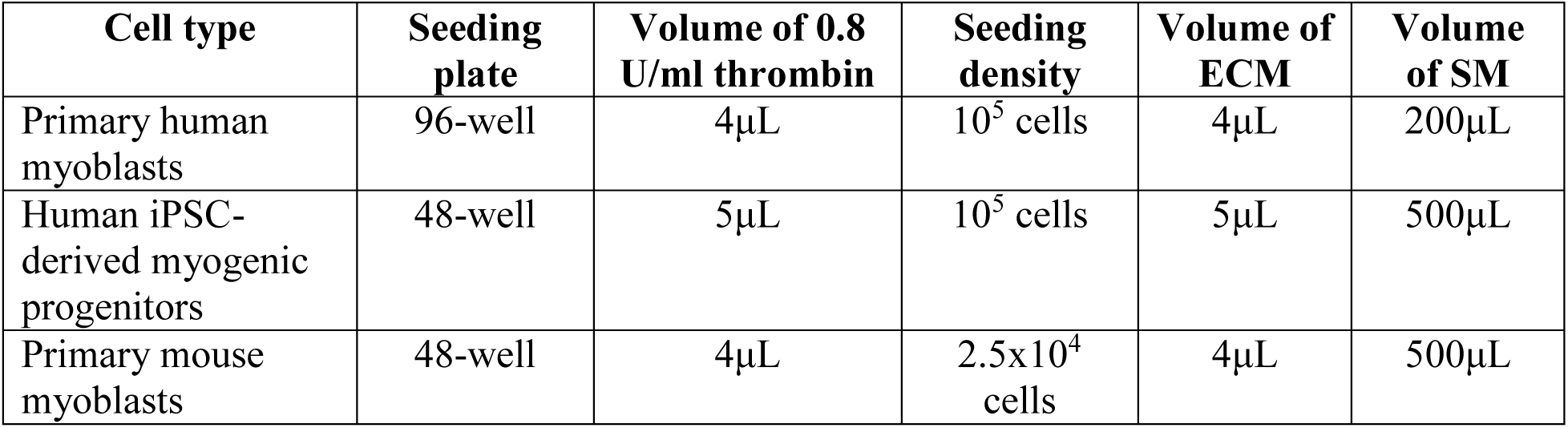
Cell densities and volumes for seeding different muscle cell lines in mini-MEndR.

### Cryopreservation of hydrogel encapsulated myoblasts seeded in paper-based scaffolds

The cryopreservation strategy aimed to closely mimic our previously established mini-MEndR workflow.[32] In mini-MEndR, scaffolds are immersed in SM immediately after seeding for 48h before changing the media to DM. Here, we strategized to “interrupt” the 48h equilibration period by freezing mini-MEndR scaffolds 24h after seeding. After thawing, cells are then cultured for the remaining 24h in SM, upon which the media is switched to DM, with half-media changes every other day thereafter.

To freeze mini-MEndR tissues, 1 mL of freezing media per tissue (90 % FBS + 10 % DMSO; Table 1) was prepared and pipetted into each cryovial (Sarstedt, #72.379.004). Using tweezers (ESD-12, anti-static stainless steel, Kaverme), scaffolds were gently picked up from their wells in the 96-well plate and deposited at the top rim of their corresponding cryovial. Once all scaffolds had been placed on the rims of cryovials, they were all successively dropped into the freezing media in the same order as they were initially picked up. This ensured that each scaffold was immersed in freezing media for a similar duration, as opposed to immediately immersing each scaffold one at a time, which would lead to greater variability between the first and last scaffolds. Cryovials were transferred to a Nalgene® Mr. Frosty (ThermoFisher, #5100-0001), and moved to a -80°C freezer where they were stored 24-48h before being transitioned to liquid nitrogen for the remainder of time (5-6 days) to complete one week of freezing.

### Thawing myoblasts cryopreserved in paper scaffolds

Prior to thawing, an empty Nalgene® Mr. Frosty was pre-chilled in a -80 °C freezer for a minimum of one hour and then moved to a Magic Ice Bucket (Grainger, Cat# WWG39C553) with the lid closed to keep the samples cool. Next, 200 μL of SM per tissue (Table 1) was prepared in a 15 mL Falcon Tube, while 225 μL pre-chilled wash media was loaded into a number of empty wells of the 96-well plate equal to the number of scaffolds that were to be thawed. A drop of wash media was added to the centre of each well that would later contain a scaffold to overcome the electrostatic forces between the tweezers and scaffold and facilitate placement of the scaffold into the well. Finally, all materials, pipettes, and reagents were calibrated and oriented in the biosafety cabinet to maximize speed during the thawing process.

Depending on user proficiency, between 4 (new handlers) to 8 (expert handlers) cryovials were removed from the liquid nitrogen tank and placed directly into the pre-chilled Nalgene® Mr. Frosty on ice. This aimed to mitigate the conflicting risks of prolonged cryovial exposure to ambient temperature and the need to repeatedly open the liquid nitrogen tank. Thawing proceeded one vial at a time, swirling the cryovial in a 37 °C water bath to dissipate excess heat and speed up thawing. Just before being fully thawed (to minimize contact time with cryoprotective agent), the vial was brought inside the biosafety cabinet and the freezing media pipetted out into a waste container. The scaffold was gently transferred to its corresponding well in the 96-well plate using tweezers, where the drop of wash media deposited earlier facilitated its placement. Next, 3 x 75 μL washes with pre-chilled SM were performed, pipetting from the 225 μL pre-set-up wells to maximize speed, and culminating with the addition of 200 μL SM. The process was repeated for each cryovial in the batch, and for each batch in the experiment (if > 8 vials). The plate was then moved to a 37 °C cell culture incubator and standard 3D culture ensued.

### Cell viability assay

Scaffolds were transferred to wells in a new plate and washed once with PBS. Staining solution was prepared by diluting Calcein AM (1:2000) and DRAQ5 (1:800) dyes in PBS as described in Table 3. 50 μL of dye solution was dispensed into each scaffold-containing well. The plate was incubated for 20 minutes in a 37 °C cell culture incubator and then imaged using confocal microscopy.

**Table 3.** Fluorescent dyes and antibodies used for immunostaining. BS=blocking solution.

| Name | Type | Dilution | Reference |
| --- | --- | --- | --- |
| Mouse anti-sarcomeric-alpha-actinin | Primary Antibody | 1:800 in BS | Sigma, #7811 |
| Mouse anti-Pax7 | Primary Antibody | 1.5:1 in PBS | Derived from DSHB cell line (Iscove) |
| Rabbit anti-Ki67 | Primary Antibody | 1:300 in BS | Abcam, #ab16667 |
| Chicken anti-GFP | Primary Antibody | 1:500 in BS | Abcam, #ab13970 |
| Goat anti-mouse AlexaFluor-488 | Secondary Antibody | 1:500 in BS | Invitrogen, #A11001 |
| Goat anti-rabbit AlexaFluor-647 | Secondary Antibody | 1:500 in BS | Life Technologies, #A21245 |
| Goat anti-chicken AlexaFluor-488 | Secondary Antibody | 1:500 in BS | Invitrogen, #A11039 |
| Phalloidin AlexaFluor-568 | F-actin probe | 1:500 in BS | Invitrogen, #A12380 |
| DAPI | Nuclear probe | 1:800 in BS | Roche, #10236276001 |
| DRAQ5 | Nuclear dye | 1:800 in BS or PBS | Cell Signalling Technology, #4084L |
| Calcein AM | Viability Dye | 1:2000 in PBS | Invitrogen, L3224 |

### Immunohistochemistry

Scaffolds were gently transferred to a new well using tweezers and washed once with PBS. Tissues were fixed with 4 % paraformaldehyde (Fisher Scientific, 50980494) for 12 min, followed by 3 x 10 min washes with PBS. For immunostaining, tissues were first blocked with 100 μL of blocking solution (Table 1) for 30 min. Then, primary antibodies were diluted in blocking solution or PBS according to Table 3, and 50 μL of the staining solution was added to each tissue. The plate was incubated overnight at 4 °C, followed by 3 x 10 min washes with PBS to wash out unbound antibody. In similar fashion, secondary antibodies were also diluted in blocking solution according to Table 3, and 50 μL of the staining solution was added to each tissue. After a 45-60 min incubation period at room temperature, the tissues were washed 3 x 10 min with PBS, and either imaged thereafter or stored in PBS at 4 °C until image acquisition.

### Image acquisition

Fixed and stained tissues were imaged using confocal microscopy (Olympus FV-1000, Olympus FluoView V4.3b imaging software) with an air objective, laser power kept less than or equal to 50 %, exposure (dwell) time 4 μs/pixel, a 1:1 aspect ratio, 640x640 box size, sequential line capture, and HV ≤ 750. HV was optimized on an image-by-image basis to maximize signal while avoiding oversaturation (red pixels on software). Scaffolds were maintained in PBS while in queue for imaging to prevent them from drying.

To evaluate tissue homogeneity, tissues were imaged at either 4X or 10X magnification, with the settings described in Table 4. SAA^+^ coverage was quantified as previously described;[27, 32] briefly, confocal image stacks were maximum Z-projected and thresholded either manually or using FIJI’s triangle algorithm, where the percentage of pixels brighter than the threshold was taken as a metric of image coverage by SAA^+^ myotubes. To assess myofiber morphology, tissues were imaged at 40X magnification, with the objective correction collar being adjusted to optimize focus (Table 4). Nuclear fusion indices were defined as the ratio between the number of nuclei present in SAA^+^ structures and the total number of nuclei in the maximum Z-projected confocal images acquired at 40X magnification. Finally, for cell viability staining, scaffolds were imaged at 20X magnification and no Kalman filtering to increase imaging speed given that the cells were not fixed (Table 4). Three to four sites were imaged from each scaffold. Cell viability was quantified as the ratio of live cells (Calcein^+^) to total cells (DRAQ5^+^) using a custom image analysis workflow (Figures S7- S9).

**Table 4.** List of imaging settings used for confocal imaging of mini-MEndR tissues with the Olympus FV-1000 confocal microscope and Olympus FluoView V4.2b imaging software.

| Imaging Parameter | 4X Images | 10X Images | 20X Images | 40X Images |
| --- | --- | --- | --- | --- |
| Purpose | Homogeneity | Homogeneity | Live/Dead | Morphology |
| Objective | 4X | 10X | 20X | 40X |
| Numerical Aperture | 0.13 | 0.3 | 0.45 | 0.6 |
| Scan Mode | Unidirectional | Unidirectional | Unidirectional | Unidirectional |
| Laser Channels | AF-488 / AF-568 / DRAQ5 (647) | AF-488 / AF-568 / DRAQ5 (647) | AF-488 / DRAQ5 (647) | AF-488 / AF-568 / DRAQ5 (647) |
| Laser Power | < 50% | < 50% | < 50% | < 50% |
| PMT Voltage (HV) | <750 | <750 | <750 | <750 |
| Amplifier Gain | 1% | 1% | 1% | 1% |
| Sequential | Line | Line | Line | Line |
| Dwell Time | 4 μs/pixel | 4 μs/pixel | 4 μs/pixel | 4 μs/pixel |
| Slice Thickness | 15-20 μm/slice | 15 μm/slice | 7-10 μm/slice | 5-8 μm/slice |
| Kalman Averaging | Line, 3 | Line, 3 | N/A | Line, 3 |
| Box Size | 640 x 640 | 640 x 640 | 640 x 640 | 640 x 640 |
| Scan Zoom | 1.0 | 1.0 | 1.0 | 1.0 |

### Calcium handling

On Day 8 of differentiation, differentiation media was removed from frozen or unfrozen mini-MEndR myotube templates. The tissues were washed 2 times with warmed PBS, and then incubated for 1 hour in the 37 °C incubator with Calbryte^TM^ 520 AM at 10 μM in Hanks buffered saline with 20 mM Hepes and 0.04 % pluronic acid. After incubation, the calcium indicator was removed, and tissues were washed twice with PBS to remove any indicator present in the well. Tissues were then returned into differentiation media and allowed to warm for 15 min in the 37 °C incubator. Calcium transients were captured in response to chemical stimulation using 2 mM acetylcholine using an inverted microscope (Olympus IX83) with a 4X magnification lens. Consecutive time-lapse epi-fluorescent images were captured at a frame speed ∼135 ms with a DP80 CCD camera (Olympus) outfitted with a fluorescein isothiocyanate filter and Olympus cellSens^TM^ imaging software. Subsequently, the fluorescent signal in the series of time-lapse images was measured using the FIJI software (ImageJ, NIH) and relative fluctuations in the fluorescence were calculated and presented as ΔF/F0 = (Fimmediate – Fbaseline)/(Fbaseline). To obtain an average ΔF/F0 transient, the transients from all technical replicates were shifted so that the time at which the ΔF/F0 first began to increase form baseline was synchronized (since the captured videos may be recording for a varying time before manually adding acetylcholine to initiate stimulation).

### Shipping myoblasts cryopreserved in cellulose scaffolds

On the day of shipping, a Styrofoam box was liberally filled with dry ice according to the expected shipping duration (4kg dry ice/day; Toronto, ON, Canada to Hamilton, ON, Canada = 4kg; Toronto, ON, Canada to Lyon, France = 12kg). Materials and reagents for cell culture and immunostaining belonging in temperatures below zero (Table 5) were parafilmed and placed in plastic bags (Falcon tubes) or an old DRAQ5 supplier container (antibodies) and placed approximately half-deep in the dry ice box. Cryovials from the liquid nitrogen were collected last (to minimize their exposure to ambient temperature) and placed in Styrofoam tube holders and a plastic bag before being placed into the dry ice box. Three cryovials containing cell-free scaffolds (paper in freezing media) were also shipped so that collaborators could practice the thawing and handling SOP without risking compromising the tissues. A temperature sensor (TempTale Dry Ice, Sensitech) was activated and placed on top of the dry ice in each box. The lid was closed and the box taped tightly. The styrofoam box was placed in a cardboard box where any extra gaps were filled with bubble wrap, a materials list (for recipients), and the materials or reagents that could be kept at room or 4 °C temperature (Table 5). Finally, the cardboard box was closed, taped, and marked with the appropriate shipping labels. Collaborators were provided our in-house protocols and all steps were performed as described in this Methods section.

**Table 5.**
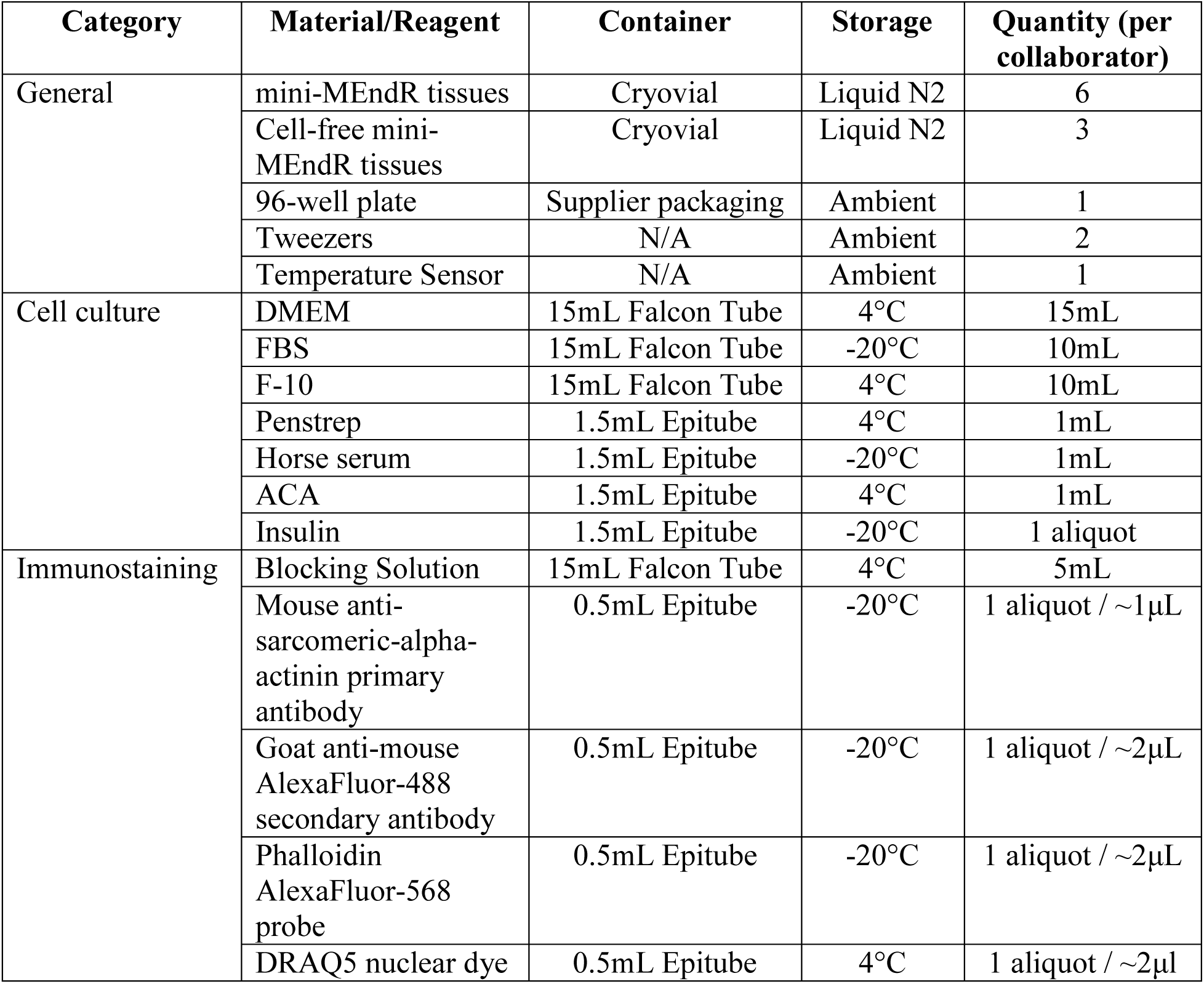
List of materials and reagents shipped to each collaborator.

| Category | Material/Reagent | Container | Storage | Quantity (per collaborator) |
| --- | --- | --- | --- | --- |
| General | mini-MEndR tissues | Cryovial | Liquid N2 | 6 |
|  | Cell-free mini-MEndR tissues | Cryovial | Liquid N2 | 3 |
|  | 96-well plate | Supplier packaging | Ambient | 1 |
|  | Tweezers | N/A | Ambient | 2 |
|  | Temperature Sensor | N/A | Ambient | 1 |
| Cell culture | DMEM | 15mL Falcon Tube | 4°C | 15mL |
|  | FBS | 15mL Falcon Tube | -20°C | 10mL |
|  | F-10 | 15mL Falcon Tube | 4°C | 10mL |
|  | Penstrep | 1.5mL Epitube | 4°C | 1mL |
|  | Horse serum | 1.5mL Epitube | -20°C | 1mL |
|  | ACA | 1.5mL Epitube | 4°C | 1mL |
|  | Insulin | 1.5mL Epitube | -20°C | 1 aliquot |
| Immunostaining | Blocking Solution | 15mL Falcon Tube | 4°C | 5mL |
|  | Mouse anti-sarcomeric-alpha-actinin primary antibody | 0.5mL Epitube | -20°C | 1 aliquot / ~1µL |
|  | Goat anti-mouse AlexaFluor-488 secondary antibody | 0.5mL Epitube | -20°C | 1 aliquot / ~2µL |
|  | Phalloidin AlexaFluor-568 probe | 0.5mL Epitube | -20°C | 1 aliquot / ~2µL |
|  | DRAQ5 nuclear dye | 0.5mL Epitube | 4°C | 1 aliquot / ~2µl |

### Image acquisition details for collaborative experiments

For the intra-provincial collaborator study, mini-myotube template tissues were mounted on a glass slide and imaged using confocal microscopy (Nikon A1R inverted confocal with a Ti2-E stand, NIS-Elements V5.4 imaging software). To evaluate tissue homogeneity, images were acquired with a 10X air objective, NA=0.45, and pixel size 0.34μm/px. To assess myotube morphology, images were acquired with a 40X water objective, NA=1.25, 0.25μm/slice, and pixel size 0.13μm/px. All images were acquired using a sequential line scan, laser power ≤ 2%, ≤ 2μs/pixel, 1:1 aspect ratio, 1024x1024 box size, and HV ≤ 100.

For the international collaborator study, mini-myotube template tissues were imaged using confocal microscopy (Zeiss LSM 880 confocal microscope, available at the CIQLE platform (SFR Santé Lyon-Est, UAR3453 CNRS, US7 Inserm, UCBL), and ZEISS ZEN 3.3 (blue edition) imaging software. To evaluate tissue homogeneity, images were acquired with a 10X air objective, NA=0.45, 15μm/slice, and HV ≤ 600. To assess myofiber morphology, images were acquired with a 40X oil objective, NA=1.4, 6μm/slice, and HV ≤ 850. All images were acquired using a unidirectional scan, averaging of 4 images, 1.64μs/pixel, 1:1 aspect ratio, and 640x640 box size.

Across all experiments, HV was optimized on an image-by-image basis to maximize signal while avoiding oversaturation (red pixels on software). Scaffolds were maintained in PBS while not being imaged to prevent them from drying. Images were provided to the Toronto-based team where they were adjusted in brightness and cropped to the same scale to account for differences in image acquisition.

### Mini-MEndR assay

Mini-myotube templates were first generated from myoblasts freshly seeded into the paper-based scaffolds, or by using myoblasts cryopreserved in paper scaffolds. All subsequent steps were performed as described in our previously optimized protocol.[32] Briefly, on Day 5 of differentiation, integrin α-7^+^ GFP^+^ muscle stem cells MACS-enriched from enzymatically digested Tg:Pax7-nEGFP (i.e. Pax7-nGFP) transgenic mouse[39] hindlimb skeletal muscle tissue were re-suspended in SAT10 media replete of FGF2. On Day 6 of differentiation, 4 μL of the re- suspended GFP^+^ muscle stem cell solution (850 cells per tissue) was dispensed onto the tissue surface, and spread evenly using a sterile cell spreader. The following day, tissues were injured with 0.5 μM cardiotoxin (CTX) diluted in differentiation medium for 4h. Tissues were then cultured for 5 days post-injury in DM supplemented with 5 ng/mL FGF2 and an inhibitor of p38 α/β MAP kinase (p38i) or DMSO carrier control. Media was completely replaced every other day. Tissues were cultured for an additional 2 days in standard DM without added compounds until fixation at 7 days post-injury (DPI).

Confocal imaging was performed using the Perkin-Elmer Operetta CLS High-Content Analysis System and the associated Harmony software as described in.[38] Images were collected using the 10X air objective (Two Peak autofocus, NA 1.0 and Binning of 1) to assess donor-derived GFP^+^ myotube coverage, and the 20X water immersion objective (Two Peak autofocus, NA 1.0 and 1.1, and Binning of 1) to enumerate muscle stem cells. Images were exported in their raw form and Max-projected and stitched using the FIJI BIOP Operetta Import Plugin (<u>ijs-Perkin Elmer</u> <u>Operetta CLS, Stitching And Export, 2022</u>).

Donor-derived cell tissue coverage (i.e. GFP^+^ signal) was quantified manually using FIJI. First, the area for analysis was demarcated using the polygon tool to crop the tissue from the entire stitched image. This permitted the exclusion of regions that appeared as empty squares in the stitched image due to focus failures encumbered during imaging. The resulting analysis area was thresholded manually and the GFP coverage recorded.

### Statistical analysis

A minimum of 2-3 independent experiments (N) each with multiple technical replicates (n) were conducted for most experiments and is detailed in Table S2. Statistical analysis was performed using GraphPad Prism 6.0 software. For comparison of two variables, statistical differences were determined by unpaired Student’s t-test or unpaired t-test with Welch’s correction. For data with more than two variables compared, a one-way ANOVA followed by Tukey’s multiple comparison test was utilized (Table S2). All values are expressed as mean ± standard deviation (SD). Significance was defined as p ≤ 0.05.

## Results

### Selection of a cellulose scaffold material for mini-myotube templates

For the mini-MEndR assay, human myoblasts are encapsulated in a fibrin/reconstituted basement membrane hydrogel mixture and seeded into rectangle-shaped cellulose-based paper scaffolds pre-adsorbed with thrombin. The myoblasts are then differentiated to form thin skeletal muscle microtissues (termed “myotube templates”) that are compatible with the footprint of 96-well culture plates (Figure 1A).[32] We first set out to identify viable paper products from readily accessible suppliers to robustly manufacture myotube templates. To this end, four paper scaffold candidates (Candidates 1-4) were preliminarily filtered from a larger pool of candidate papers based on thickness and material (cellulose) similarity to the original paper scaffold used to produce mini-myotube templates that was no longer commercially available. Scanning electron microscopy (SEM) images indicated that Candidate 1 and Candidate 3 shared a similar porosity and morphology of paper fibers to the original cellulose paper, while Candidate 2 and Candidate 4 exhibited a much more compact microstructure (Figure S1). Indeed, quantification of the image porosity confirmed that Candidate 1 and Candidate 3 had the highest pore area coverage and a pore area closest to the original cellulose paper (Figure S2). However, further analyses revealed that Candidate 1 had a much higher number of smaller sized pores as compared to the original candidate (Figure S2). Candidate 4 was omitted from testing at this stage given its greatest disparity from the original candidate across all metrics (Figures S1-S2).

**Figure 1.**
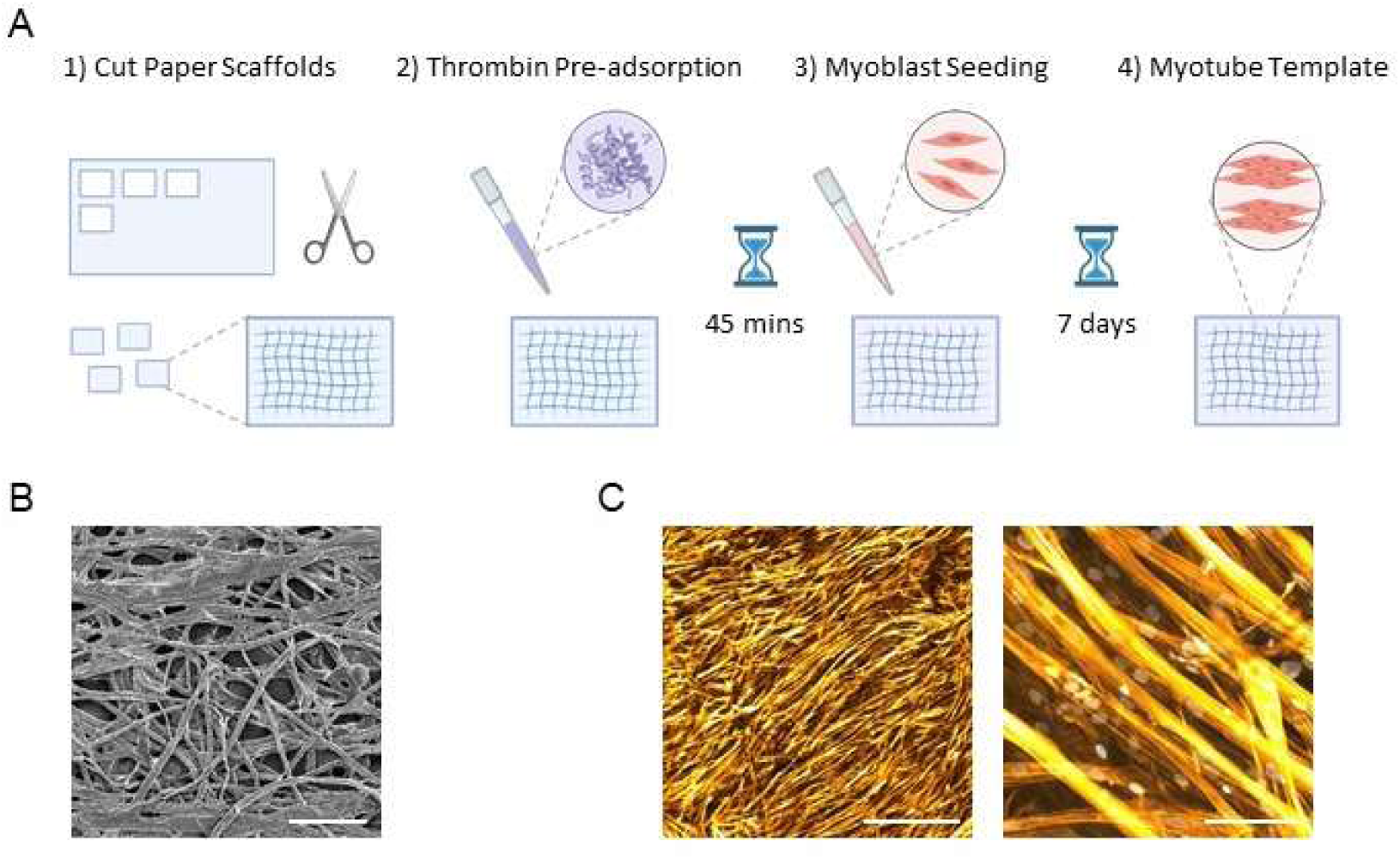
Formation of skeletal muscle miniaturized myotube templates in cellulose scaffolds. **(A)** Workflow to produce miniaturized skeletal muscle myotube templates. On Day 0, primary human myoblasts were seeded onto rectangular cellulose-based paper scaffolds pre-adsorbed with thrombin and cultured in SM in a 96-well plate. On Day 2, the culture medium was switched to differentiation medium and myoblasts are differentiated over 7 days, generating a thin layer of 3D scaffold-supported human muscle microtissues (“myotube template”). **(B)** Cropped 800μm x 800μm section from a representative 50X scanning electron microscopy (SEM) image of the paper material selected to serve as the scaffold for myotube templates (Candidate 1). Scale bar = 200μm. **(C)** Representative 4X and 40X confocal images of primary human myoblasts differentiated in mini-MEndR over 7 days. Actin is shown in orange (Phalloidin), and nuclei are shown in grey (DRAQ5). 4X scale bar = 1000μm; 40X scale bar = 100μm.

We next evaluated the ability of each paper candidate to support the generation of myotube templates that were comparable to those generated in the original paper scaffold. To do this, we seeded primary human myoblasts (Figure S3), primary mouse myoblasts (Figure S4), and human iPSC-derived myogenic progenitors (Figure S5) within the candidate papers and differentiated them for up to 2-weeks. In all candidate papers, the myoblasts differentiated to form elongated, multinucleated myotubes that were locally aligned but globally disorganized, as observed in the original cellulose scaffold (Figures S3-S5). However, only Candidate 1 supported a total myotube area coverage that was comparable to the original paper-based scaffold across all timepoints and cell types tested (Figures S3-S5). Therefore, Candidate 1 was selected as the paper scaffold material for all subsequent studies (Figure 1B-C).

### Development of a protocol for cryopreserving myoblasts in paper-scaffolds

We next set out to develop a protocol to cryopreserve myoblasts encapsulated in hydrogel and seeded into paper-based scaffolds with a view to enabling broad uptake of the mini-MEndR platform within the scientific community. We based our protocol on previous work performed by Grant and colleagues[33] in which myoblasts were encapsulated in a hydrogel mixture of 30 % Matrigel^TM^, 4 mg/mL fibrinogen, and 0.5 U/mg-fibrinogen thrombin, and seeded into polyethylene glycol dimethacrylate (PEGDMA) ring-shaped molds. The rings were cryopreserved either before myoblast differentiation (ECM containing mononucleated myoblasts) or after 7 days of differentiation (ECM containing multinucleated myotubes). Importantly, only the cryopreserved undifferentiated myoblasts, had the capacity to acquire expression of mature myogenic markers (e.g. MHC-1, MYF6) and produce aligned myotubes alignment upon thawing and differentiating. [33] We therefore decided to cryopreserve myoblasts in the paper-based scaffold in the undifferentiated state.

For our cryopreservation method, myoblasts were encapsulated in a fibrin-based hydrogel and seeded into the cellulose paper exactly as described in the original mini-MEndR assay (Figure 1A). Based on iterative research and pilot work (see Table S1 and Figure S6), we established a cell seeding density and cryopreservation regimen that maximized cell viability. In brief, myoblasts are seeded and cultured in Seeding Medium (SM) for 48h before transitioning to Differentiation Medium (DM) in the original protocol used to generate mini-myotube templates. We converged on cryopreserving myoblasts 24h post-seeding, a timepoint halfway through this SM equilibration step. Thus, after 24h in SM, cell-laden scaffolds were transferred to commercial cryovials containing freezing medium (90 % FBS + 10 % DMSO) and slow cooled for 24-48h in a Nalgene® Mr. Frosty in a -80 °C freezer (Figure S6). Cryovials were then transferred to a Dewar filled with liquid nitrogen for the remaining time (5-6 days) to complete one week total of freezing. After storage, cryovials were quickly warmed in a water bath and the scaffolds were transferred to wells of a 96-well plate for continued culture (Figure 2A).

**Figure 2.**
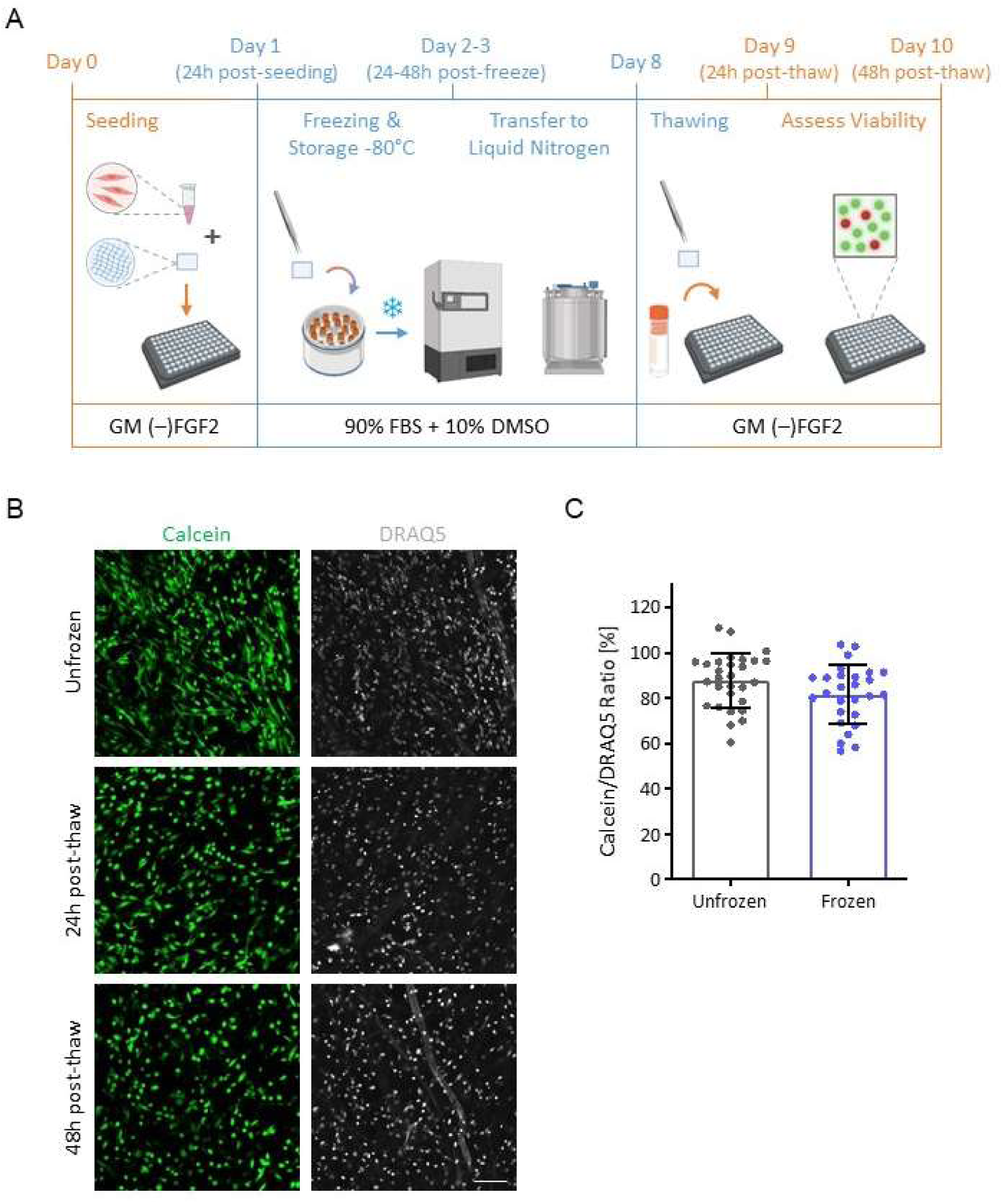
Cell viability of primary human myoblasts after cryopreservation in cellulose paper scaffolds. **(A)** Schematic of experimental procedure and culture medium timeline for assessing the viability of myoblasts cryopreserved in cellulose scaffolds. After reaching confluency in 2D culture, myoblasts are seeded in paper scaffolds in a 96-well plate. After 24h, scaffolds are transferred to cryovials containing freezing medium (90% FBS, 10% DMSO) and placed in a Nalgene® Mr. Frosty in a -80°C freezer for slow cooling. Between 24-48h later, cryovials are transferred to a liquid nitrogen dewar and stored there for the remainder of the 7 days of total freezing. Scaffolds are then extracted from the cryovials during thawing and cultured in 96-well plates. Cell viability is assessed by a live/total assay at both 24h and 48h post-thawing. GM (–) FGF2 = growth media without fibroblast growth factor 2 (Table 1). **(B)** Representative 20X live/total confocal images of primary human myoblasts in cellulose scaffolds either unfrozen, 24h post-thaw, or 48h post-thaw. Scale bar = 100μm. Live cells are shown in green (Calcein) and cell nuclei are show in grey (DRAQ5). The dead channel (propidium iodide) was omitted due to it staining the paper fibers. **(C)** Quantification of cell viability (live cells / total cells). Cells were counted using a custom developed algorithm in FIJI (Figures S7-S9). Each plotted point represents an image site. Statistics performed using unpaired two-tailed t-test, ns – no significance. N=4 biological replicates, n=2-3 technical replicates (scaffolds), with s=3-4 sites each.

To assess the effectiveness of our cryopreservation workflow we quantified cell viability 24h and 48h post-thawing by analyzing staining for Calcein and DRAQ5 using confocal imaging. Cell viability was quantified as the ratio of live cells (Calcein^+^) to total cells (DRAQ5^+^) using a custom image analysis workflow (Figures S7-S9). Representative confocal images revealed that myoblasts remained viable at all timepoints post-thaw and that they maintained a homogenous distribution across the scaffold (Figure 2B). We observed that cell viability in unfrozen (tissues that were never frozen) tissues was 87.8% ± 8.1%, while the viability of the frozen myoblasts was slightly lower at 81.6% ± 11.7% (Figure 2C). We also observed that cells in the frozen tissues had a slightly more rounded morphology relative to their unfrozen counterparts (Figure 2B). We therefore concluded that while the cells were viable after thawing it was important to determine whether the observed morphological differences would influence myoblast maturation or function at later timepoints in culture.

### Myoblasts cryopreserved in paper-scaffolds form myotube templates with unperturbed morphology and calcium release kinetics

We next evaluated whether myoblasts cryopreserved in the cellulose scaffolds would retain the ability to differentiate into myotube templates comparable to unfrozen controls. In this experiment, after 1-week of cryopreservation, frozen tissues were thawed into SM for 24h, and then switched to DM for 7 days (Figure 3A).

**Figure 3.**
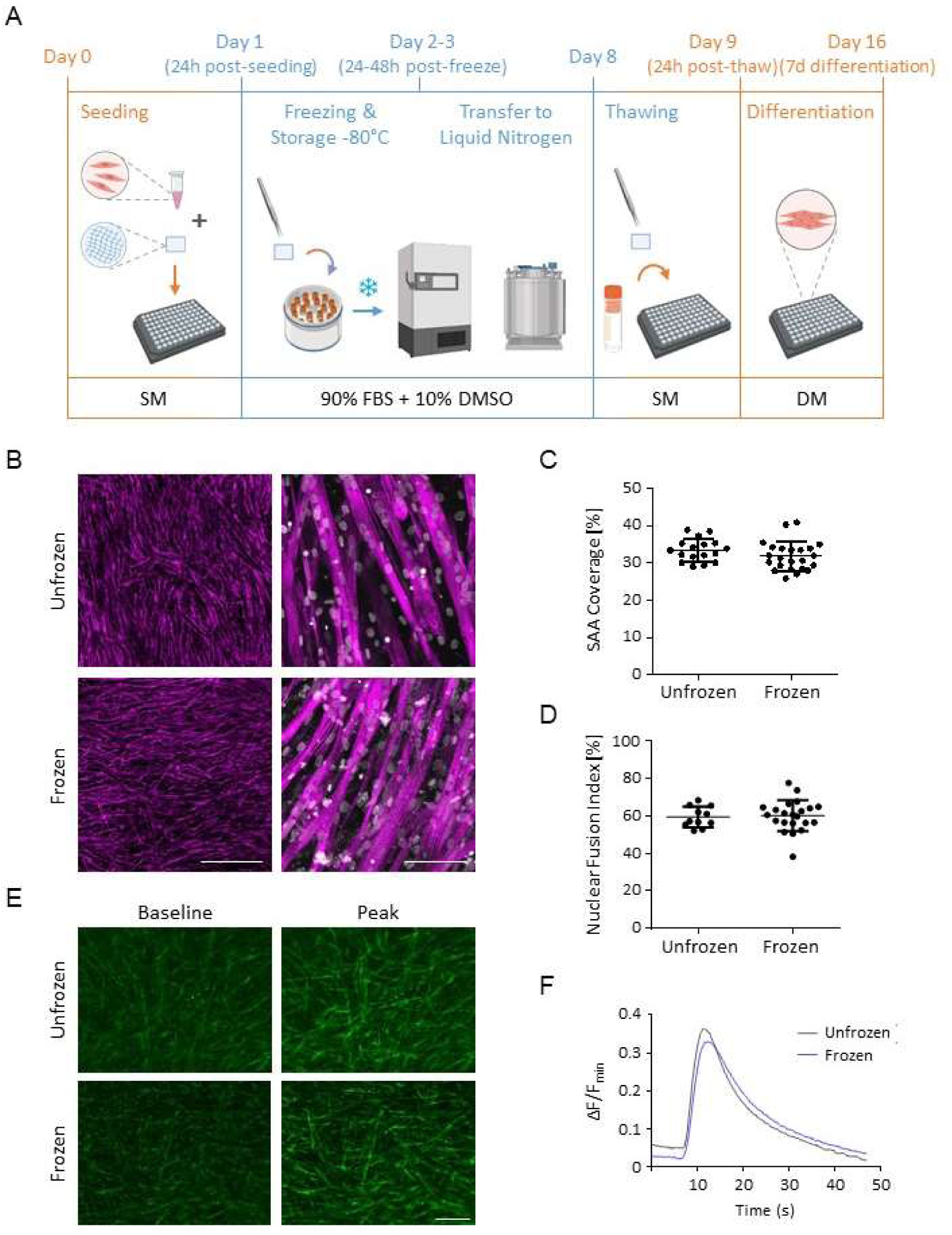
Evaluation of differentiation of primary human myoblasts after cryopreservation in cellulose scaffolds. **(A)** Schematic of experimental procedure and culture medium timeline for assessing the differentiation of myoblasts cryopreserved in paper scaffolds. After reaching confluency in 2D culture, myoblasts were seeded in paper scaffolds in a 96-well plate. After 24h, scaffolds were transferred to cryovials containing freezing medium (90% FBS, 10% DMSO) and placed in a Nalgene® Mr. Frosty in a -80°C freezer for slow cooling. Between 24-48h later, cryovials were transferred to a liquid nitrogen dewar and stored there for the remainder of the 7 days or three months of total freezing. Scaffolds were then extracted from the cryovials during thawing and cultured in 96-well plates. After a 24h equilibration period and 7 days of differentiation, tissues were fixed and immunostained to assess myotube coverage and morphology or stimulated with Acetylcholine to assess calcium transients. SM = Seeding Medium (Table 1). **(B)** Representative 4X and 40X confocal images of tissues after 7 days of differentiation. Staining of sarcomeric-alpha-actinin (SAA) is shown in magenta, and nuclei are shown in grey (DRAQ5). 4X scale bar = 1000μm; 40X scale bar = 100μm. **(C)** Quantification of sarcomeric-alpha-actinin (SAA) fiber area coverage between frozen and unfrozen (never frozen) tissues. Statistics performed using unpaired two-tailed t-test, ns – no significance. N=5 biological replicates, n=2-5 technical replicates (scaffolds). **(D)** Quantification of nuclear fusion index between frozen and unfrozen tissues. Statistics performed using unpaired two-tailed t-test, ns – no significance. N=3 biological replicates, n=3-6 technical replicates (scaffolds), with s=1-2 sites each. **(E)** Representative 4X fluorescence images at baseline and peak fluorescence and **(F)** calcium transients of unfrozen and frozen tissues. Primary human myoblasts differentiated over 8 days were incubated with Calbryte^TM^ 520 calcium indicator dye and stimulated with 2mM Acetylcholine. Fluorescence intensity over time was normalized by taking the difference between the immediate and baseline fluorescence intensities and dividing by the baseline intensity ((Fimmediate – Fbaseline)/Fbaseline). Scale bar = 200μm. Curve represents average of N=1 experimental replicate, n=5-6 technical replicates (scaffolds). See Figure S10 for additional characterizations and explanation of metrics.

Using fluorescent confocal microscopy, we found that cryopreserved myoblasts differentiated into elongated myotubes with homogenous coverage of the tissue (Figure 3B). When we quantified myotube area coverage, we observed comparable coverage between the frozen and unfrozen control; with a slight but non-significant reduction in the mean sarcomeric-alpha-actinin (SAA) coverage among frozen tissues (33.4% ± 3.0% to 31.8% ± 3.9% respectively) (Figure 3C). Higher magnification images showed that cryopreserved myoblasts formed myotubes that were elongated, striated, and locally aligned (Figure 3B), and quantification affirmed similar nuclear fusion indices between unfrozen and frozen tissues (59.3% ± 5.5% and 60.0% ± 8.2% respectively; Figure 3D). Together, this data suggested that cryopreservation did not significantly impact the differentiation potential of myoblasts in the paper-based scaffolds.

Given the encouraging morphological results, we next performed functional analysis of the myotubes by assessing their capacity to mobilize calcium in response to an acetylcholine (ACh) stimulus. Briefly, myotube template tissues on Day 8 of differentiation were incubated with Calbryte^TM^ 520 AM calcium indicator dye and subsequently stimulated with 2 mM ACh. Time-lapse calcium transients were captured at 4X magnification (Supplementary Videos S1 and S2). Both unfrozen and frozen tissues demonstrated the expected increase in fluorescence intensity upon stimulation, affirming that the myotubes in both conditions release calcium upon ACh stimulation (Figure 3E). Tissues derived from myoblasts cryopreserved in cellulose scaffolds also displayed similar transients to unfrozen tissues, including a peak ΔF/F0 of 0.33, a non-statistically significant drop from the 0.36 average produced by unfrozen controls (Figure 3F). To more closely analyze the transients, we computed the metrics of full-width at half-maximum (FWHM), time to peak (TTP), and rate of calcium release (slope of ΔF/F0 immediately after stimulation), explained in Figure S10A. Only TTP was statistically significant different between frozen and unfrozen tissues (Figure S10E). However, a trend of slightly delayed calcium responses by frozen tissues was observed across both other metrics (Figure S10B-D). Taken together, our data suggest that myoblasts cryopreserved in the paper-based scaffold differentiate into morphologically comparable and ACh-responsive myotube templates, positioning the assay well for facilitating inter-laboratory studies of skeletal muscle health.

### Cryopreservation in paper-based scaffolds is amenable to different myoblast sources

We next assessed whether our cryopreservation SOP was compatible with myotube templates derived from another muscle cell type used by our group. Specifically, human induced-pluripotent stem cell (iPSC)-derived myogenic progenitors were encapsulated in hydrogel, seeded into the paper-based scaffold and subjected to the previously described cryopreservation protocol (Figure 3A). The thawed and differentiated cellulose scaffolds seeded with iPSC-derived myogenic progenitors generated homogeneous myotube templates that were locally aligned and globally disorganized (similar to unfrozen controls) (Figure 4A). Quantification of SAA coverage demonstrated no significant differences between unfrozen and frozen tissues (37.5% ± 2.7% and 37.4% ± 4.3% respectively; Figure 4B). Furthermore, higher magnification images revealed that myotubes were elongated, striated, and multinucleated and qualitatively thicker in diameter when compared to the myotubes in the unfrozen control (Figure 4A). This data suggested that our protocol to cryopreserve mini-MEndR tissues can be extended to other sources of skeletal muscle myoblasts, making it a versatile system for 3D myotube studies.

**Figure 4.**
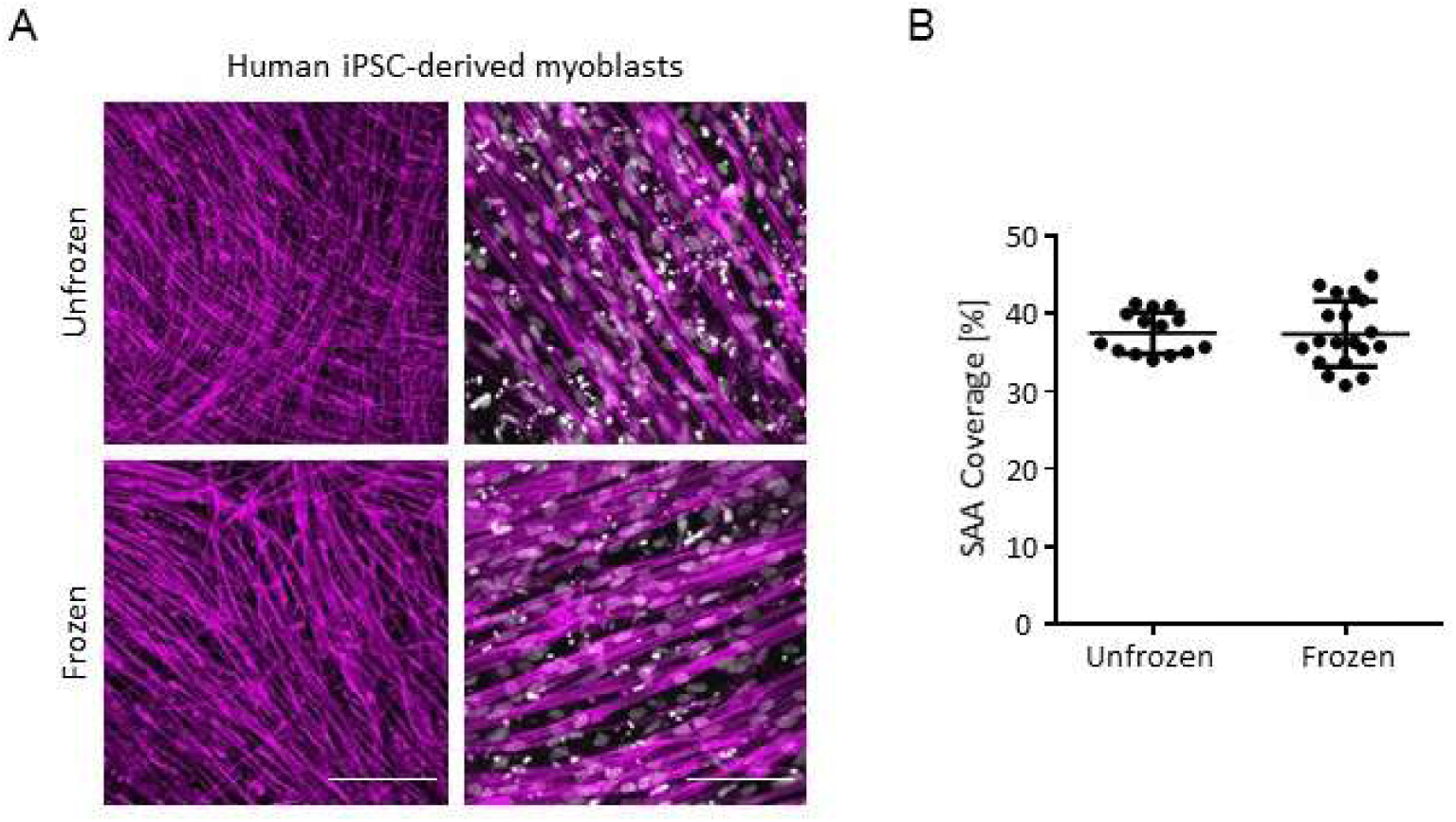
Cryopreservation protocol is amenable to human iPSC-derived myogenic progenitors. **(A)** Representative 10X and 40X confocal images of human iPSC-derived myogenic progenitors either unfrozen (never frozen) or frozen in cellulose scaffolds and cultured over 7 days of differentiation. 10X scale bar = 400μm; 40X scale bar = 100μm. Sarcomeric-alpha-actinin (SAA) is shown in magenta, and nuclei are shown in grey (DRAQ5). **(B)** Quantification of SAA coverage from 10X images of human iPSC-derived myogenic progenitors either unfrozen or frozen in cellulose scaffolds and cultured over 7 days of differentiation. Statistics performed using unpaired two-tailed t-test, ns – no significance. N=2 biological replicates, n=4-6 technical replicates (scaffolds).

### Cryopreservation in paper-based scaffolds facilitates inter-laboratory collaborative studies of skeletal muscle biology

To demonstrate proof of concept that our cryopreservation protocol enables the adoption of 3D muscle biology by non-experts in the scientific community, we assessed the impact of shipping myoblasts cryopreserved in paper-based scaffolds. We shipped cryopreserved myoblast template tissues to domestic and international collaborators and then compared the quality of muscle tissues they produced upon thawing and then differentiating the tissues in culture. For this, a batch of primary human myoblasts was used to seed numerous cellulose scaffolds, which were transferred to cryovials, stored in liquid nitrogen for 2 months and then distributed (Figure 5A) for parallel studies in-house, and at a domestic (McMaster University (Hamilton, ON, Canada)) and an international site (Institut Neuromyogène (Lyon, France)).

**Figure 5.**
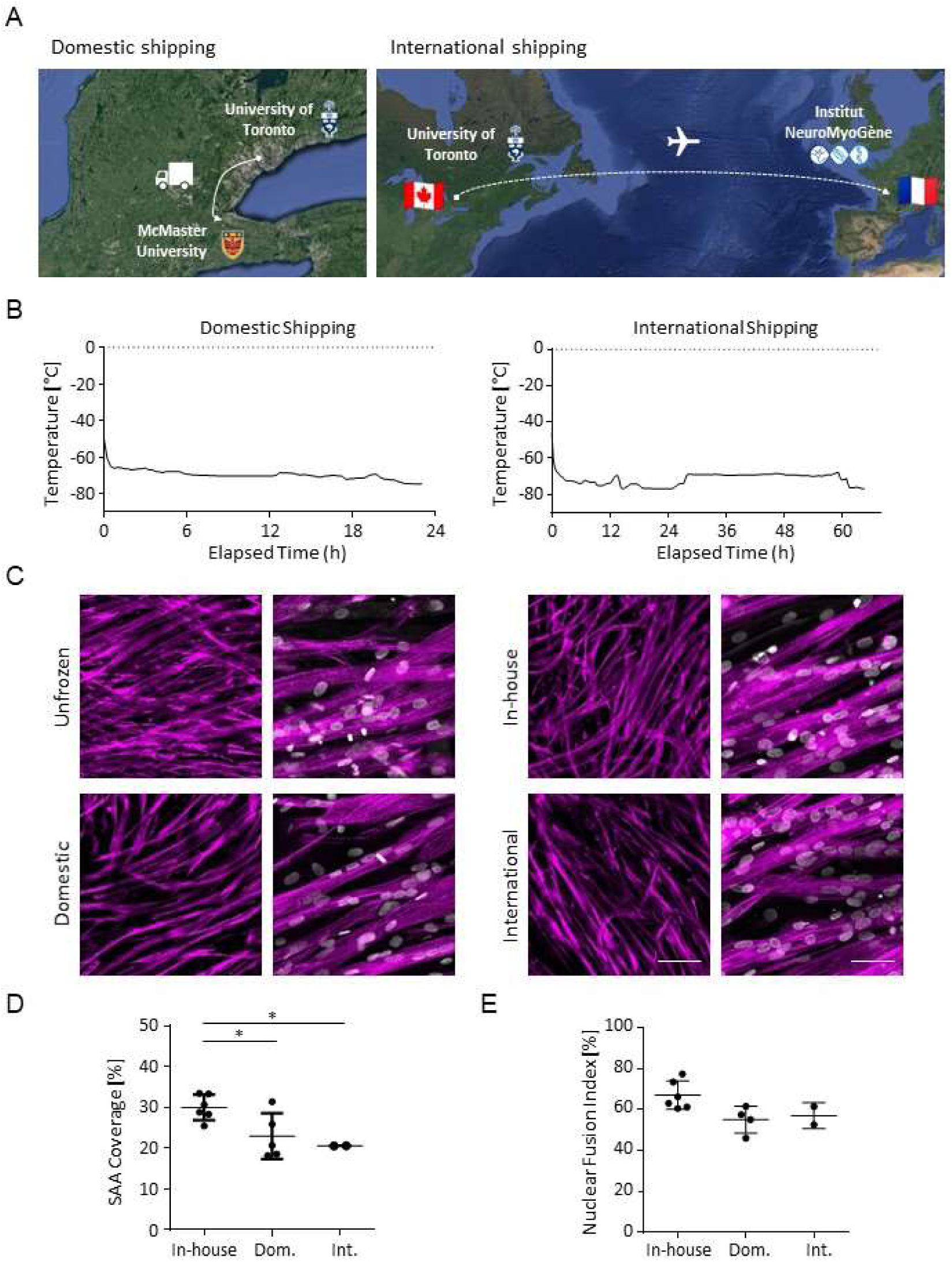
Cryopreservation in cellulose scaffolds enables inter-laboratory collaborative studies of skeletal muscle biology. **(A)** Satellite map of institutions involved in collaborative study. Myoblasts cryopreserved in cellulose scaffolds were shipped domestically via ground between the University of Toronto (Toronto, ON, Canada) and McMaster University (Hamilton, ON, Canada) and internationally via air between the University of Toronto (Toronto, ON, Canada) and the Institut NeuroMyogène (Lyon, France). **(B)** Temperature profiles during shipment. Temperature was recorded at 1-minute intervals using the Sensitech TempTale Dry Ice temperature sensor. Customized graphs were created by manually interpolating datapoints from the sensor’s automatically generated temperature graphs and inputting them into the PRISM graphing software. **(C)** Sample 10X and 40X confocal images of primary human myoblasts either (i) unfrozen or (ii-iv) frozen for 2 months and shipped either (ii) in-house, (iii) domestically, or (iv) internationally, and subsequently differentiated for 7 days. Sarcomeric-alpha-actinin (SAA) is shown in magenta, and nuclei are shown in grey (DRAQ5). 10X scale bar = 200μm; 40X scale bar = 50μm. **(D)** Quantification of SAA coverage from 10X images of in-house, domestic, and internationally shipped tissues. Statistics performed using one-way ANOVA with Tukey post-test, *p<0.05. N=1 biological replicates, n=2-6 technical replicates (scaffolds), averaged from s=1-3 sites each. **(E)** Quantification of nuclear fusion index from 40X images of in-house, domestic, and internationally shipped tissues. Statistics performed using one-way ANOVA with Tukey post-test, ns – no significance. N=1 biological replicates, n=2-6 technical replicates (scaffolds), averaged from s=2- 4 sites each.

Temperature monitoring devices (TempTale Dry Ice, Sensitech) were packed into each shipping container to record the temperature at 1-minute intervals throughout the shipping process. Temperature predominantly held steady between -65°C to -75°C during shipping (Figure 5B). By assessing in-house controls (unfrozen and cryopreserved), we confirmed that the selected cell batch formed differentiated myotube templates with homogenous coverage and that myotubes were also locally aligned, striated, and multinucleated (Figure 5C). Through this 2-month cryopreservation study (Figure 5) and a subsequent 3-month cryopreservation study (Figure S11), we validate the fidelity of myoblast tissues following long-term cryopreservation.

The cryopreserved samples processed by the domestic and international collaborators also generated differentiated myotubes that were elongated and multinucleated, with nuclear fusion indices comparable to in-house controls and within the range we previously reported for myotube templates (in-house: 67.1% ± 7.0%, domestic: 55.1% ± 6.6%, international: 57.1% ± 6.3%) (Figure 5E).[27] However, quantification of SAA coverage suggested that myotube templates derived from both domestic and international shipping were lower than in-house controls, with the former having slightly higher coverage than the latter (in-house: 30.0% ± 3.1%, domestic: 22.9% ± 5.6%, international: 20.6% ± 0.3%) (Figure 5D). This was consistent with our observations in-house that some user experience with the thawing and handling steps may be required to attain full proficiency with the platform (Figure S12). Nevertheless, this data show that our cryopreservation technique permits long-term (multi-month) storage, which in turn allows for flexible assay provision and inter-laboratory collaborations.

### Myoblasts cryopreserved in cellulose-paper can be used to conduct studies of muscle endogenous repair (MEndR)

We next assessed whether cryopreservation had any impact on the capacity of the myotube template to enable muscle stem cell engraftment and subsequent injury and repair of the muscle template (the MEndR repair assay). To do this, we performed a complete MEndR repair assay.[27, 32] Briefly, we generated myotube templates derived from myoblasts either unfrozen or cryopreserved in mini-MEndR (Figure 3A) and engrafted GFP^+^ murine muscle stem cells on Day 6 (Figure 6A). The following day, we injured the myotube template with cardiotoxin (CTX), a snake venom toxin, leading to a reduction in myotube template fiber area coverage by ∼50% relative to uninjured (CTX-) controls as optimized by our previous protocol[32] (Figure 6A-B). We then treated a subset of the injured tissues with either a vehicle control or an inhibitor of p38 α/β MAP kinase (p38i), a known stimulator of murine muscle stem cell self-renewal division in culture[7, 27–30] (Figure 6A). Tissues were then cultured for 7 days to allow for stem cell mediated repair of the injured muscle tissue. After 7 days of culture post-injury, we observed no statistically significant differences in the stem-cell mediated repair of unfrozen versus cryopreserved tissues (Figure 6C). Further, as expected p38i-treated tissues exhibited significantly higher donor (GFP^+^) myotube coverage area relative to DMSO carrier control in both unfrozen and frozen myoblast-derived myotube templates (unfrozen: DMSO CTX+ 10.2% ± 1.3%, CTX+ p38i 14.9% ± 4.0%; frozen: DMSO CTX+ 10.9% ± 2.6%, CTX+ p38i 14.1% ± 2.3%) (Figure 6C-D). This data suggests that cryopreserving myoblasts in a paper-based scaffold for eventual use to produce the myotube templates that form the basis of the mini-MEndR assay is a feasible strategy to facilitate adoption of the MEndR assay by the broader scientific community.

**Figure 6.**
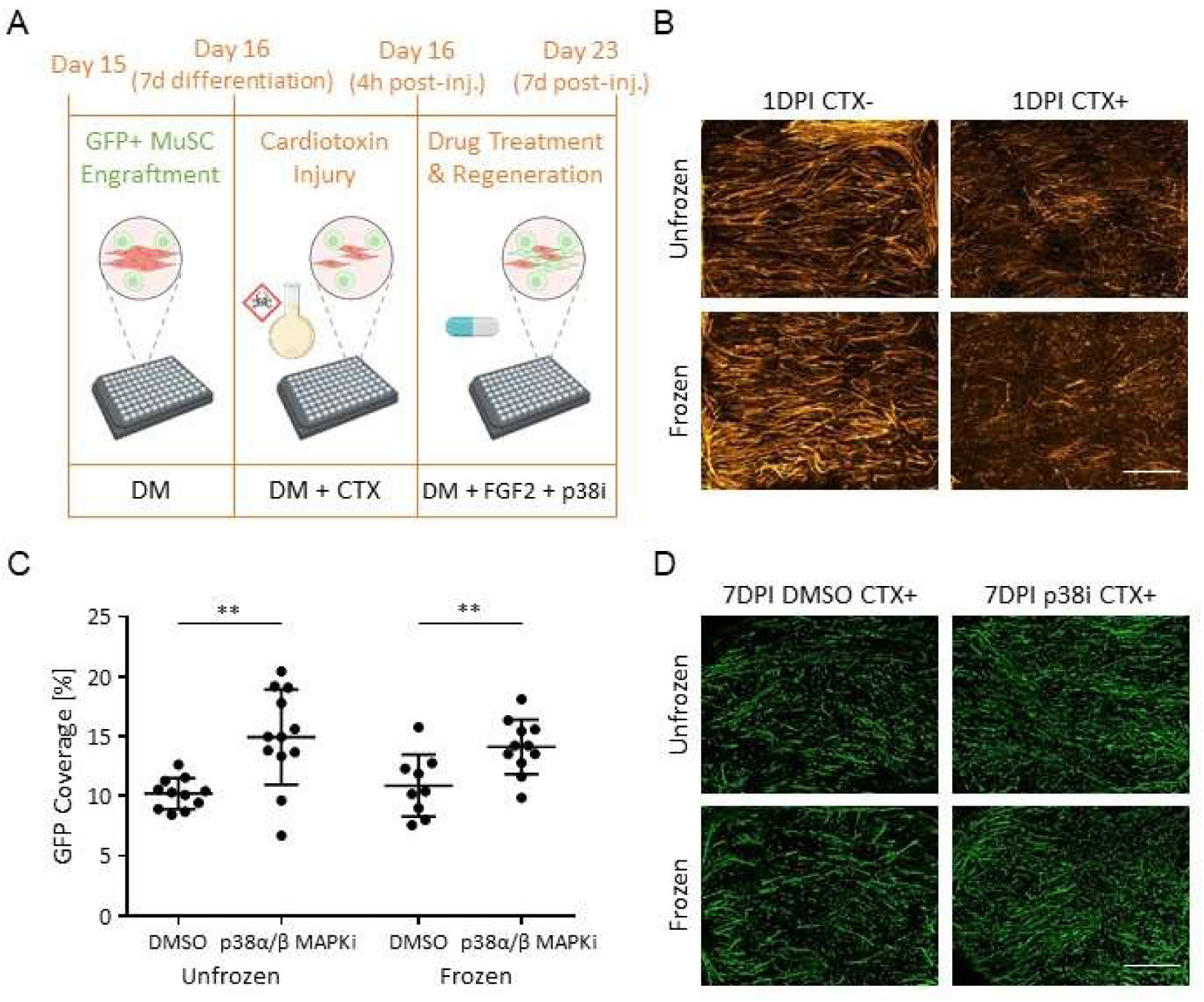
Muscle stem cells engrafted onto myotube templates derived from primary human myoblasts cryopreserved in cellulose scaffolds recapitulate expected responses to a known stimulator of muscle stem cell-mediated muscle tissue repair. **(A)** Schematic of experimental procedure and culture medium timeline for MEndR injury and regeneration assay, consisting of first the generation of myotube templates derived from myoblasts cryopreserved in cellulose scaffolds (Figure 3A), then engraftment of GFP^+^ murine muscle stem cells followed by cardiotoxin injury and muscle stem cell-mediated regeneration with either p38i or DMSO carrier control. **(B)** Representative 10X stitched confocal images of unfrozen or frozen-derived myotube templates 1- day post-injury (DPI). Actin is shown in orange (Phalloidin). 10X scale bar = 1000μm. **(C)** Quantification of GFP coverage and **(D)** representative 10X stitched confocal images of muscle stem cell-mediated repair of unfrozen or frozen-derived myotube templates with either p38i or DMSO carrier control at 7DPI. Donor-derived myotubes are shown in green (GFP). 10X scale bar = 1000μm. Statistics performed using two-tailed t-test between p38i-treated and control for each of unfrozen and frozen datasets, **p<0.01. N=3 biological replicates, n=3-5 technical replicates (scaffolds) each.

## Discussion

In vitro 3D culture models provide an exciting opportunity for studying human skeletal muscle biology at a throughput higher than is possible in animal models. Adoption of these culture models however is hindered by the requirement of in-house manufacturing expertise and specialized instruments. To increase the potential of model adoption of a culture model established by our team (the mini-MEndR platform), we established a cryopreservation protocol to enable storage and transportation of 3D myoblast tissues following in-house manufacturing. Our approach increases the accessibility of our culture platform, in particularly for groups with less myoblast culture or seeding expertise, or those seeking an off-the-shelf, validated assay to study skeletal muscle biology.

The two most common methods for cell and tissue freezing are slow cooling and vitrification. Vitrification requires extremely rapid cooling rates to bypass ice nucleation and crystallization, and is typically restricted to samples with very small volumes (microliters) that can tolerate high concentrations of cryoprotective agent.[40] Thus, we employed the slow cooling method (- 1°C/min), which is less technically challenging, subject to less variability, and has been previously employed with myoblasts.[33] Initial pilot work, building on Grant et al.,[33] used a cryopreservation freezing medium widely used in the field[41–43] of 10% DMSO and 90% FBS, the latter which is believed to support cryopreservation by stabilizing the cell membrane, decreasing extracellular ice formation, preventing excessive concentration of solutes inside the cell, and minimizing cell dehydration.[44–45] Our pilot experiments also highlighted the importance of certain specific steps in our methods that were critical to protocol success summarized in Figure S6. Specifically we realized it was important to (i) keep the freezing medium cool given the temperature-dependent cytotoxicity of DMSO,[46–49] (ii) immerse all tissues into freezing medium as quickly in succession as possible to minimize replicate-to-replicate variation, (iii) begin the slow cooling process immediately after scaffolds had been immersed in freezing medium, (iv) thaw cryovials as quickly as possible, and remove scaffolds from the freezing medium instantly thereafter, (v) perform washes with wash media to scaffolds once removed from the freezing medium, and (vi) keep cryovials frozen until immediately before starting the thaw process. For example, this meant positioning all scaffolds along the rim of cryovials to subsequently immerse them into freezing medium simultaneously (for (ii)), and keeping the cryovials in a pre-frozen Nalgene® Mr. Frosty and ice bucket until right before thawing (for (vi)). We also limited freezing to a single scaffold per cryovial to enable downstream correlation between tissue viability and/or morphology to the freezing/thawing order. This allowed any issues with the protocol to be easily identified.

A significant difference between the MEndR template and the format used to cryopreserve myoblast tissues in previous work[33] is the presence of the cellulose scaffold. We rationalized that the thin geometry of the scaffold sheet, and resulting thin myotube template was likely beneficial to the cryopreservation process to facilitate the rapid temperature changes necessary during the thaw step to avoid ice crystal formation and compromised cell viability. We did have concerns however, that the cryoprotective agent (DMSO) might absorb into the paper fibres and negatively impact long-term cell viability. Our analysis of post-thaw cell viability suggested however, that cell viability was not dramatically different between unfrozen (never frozen) (87.8% ± 8.1%) and thawed samples (81.6% ± 11.7%). Further, some cell death due to cryopreservation was expected[50] and was also observed by Grant et al. who achieved 93% ± 8.4% myoblast viability for thawed samples relative to never frozen samples.[33] In comparison the viability achieved after thawing using our protocol with the cellulose scaffold present was similar (93% ± 14.7% relative to the unfrozen samples). This suggests that the cellulose scaffold did not appear to interact with the DMSO solvent in any obviously detrimental ways. Interestingly, we did observe slight morphological differences between unfrozen and post-thawed myoblasts. Specifically, never frozen myoblasts took on an elongated morphology while post-thawed myoblasts were more spherical. This is consistent with previous reports in which cryopreserved primary human myoblasts exhibited a spherical morphology after thawing,[51] suggesting that this effect was likely not due to the format of the mini-MEndR scaffold specifically.

Despite the varying morphologies at early timepoints, we observed that cryopreserved myoblasts differentiated to form homogeneous myotube templates with comparable SAA coverage to unfrozen controls (31.8% to 33.4% respectively). Given that we also observed a slight decrease in cell viability among frozen tissues, the slight decrease in SAA coverage could potentially be attributed to this slight reduction in viable cell density. SAA coverage was also much less variable between tissues compared to the viability data, suggesting that differentiating myoblasts perhaps reach a plateau in SAA coverage. Morphologically, both unfrozen and post-thawed myoblasts formed elongated, multi-nucleated and striated myotubes. These myotubes were also locally aligned and globally disorganized, as observed by Davoudi et al. in the original MEndR platform.[27] Calcium responses were similar in myotube templates produced from myoblasts immediately after seeding into the paper-based scaffold or after a period of cryopreservation. Further, muscle stem cells were capable of engrafting into the myotube templates produced from cryopreserved myoblast tissues, and upon injury produced stem-cell mediated muscle template repair similar to unfrozen templates. Both morphologically and functionally therefore, the cryopreservation protocol, and any interactions between the DMSO solvent and cellulose fibres did not appear to significantly impact the properties of differentiated myotube MEndR templates.

As proof-of-concept that cryopreserved myoblast template tissues could logistically be shipped, thawed and used by collaborators in another institution, we shipped cryopreserved myoblast templates to local and international collaborators. We packed cell-free scaffolds (papers in freezing media only) to provide collaborators a low-risk opportunity to practice thawing and handling. Further, we included temperature monitoring devices to enable us to validate that samples remained frozen throughout the shipping process. Samples were shipped via truck (domestically) or via air (internationally) by FedEx. We observed slightly reduced SAA coverage in differentiated myoblast templates that were thawed, differentiated, and analyzed by collaborators. We speculate that these slight differences were most likely due to collaborators having limited experience with 3D myoblast tissue cultures and sample processing for analysis. Alternatively, while temperature profiles indicated suitable temperature control during the shipping process, it is possible that exposure to ambient conditions for an unlogged duration could have occurred during the unpacking or transferring of cryovials. We note that collaborators did not use thawed myoblast template tissues to conduct a complete stem-cell mediated repair assay as isolation of the muscle stem cells required for engraftment is technically challenging and requires significant optimization. We recognize this is a limitation of our proposed cryopreservation approach. Nonetheless, the results from our in-house endogenous repair assay with thawed myoblasts coupled with the successful generation of elongated, multi-nucleated myotube templates after shipping will make the full MEndR assay more feasible for users. Furthermore, for researchers with technical or experimental design barriers for stem cell engraftment, the generation of easy-to-manufacture and easy-to-analyze myotube templates alone facilitates the study of many important questions from myotube biology, toxicity screens on myotube health, and the study of mono-nucleated Pax7^+^ “reserve” cells that are present in the myotube template.[32]

## Conclusion

We have developed a protocol for the cryopreservation of myoblasts in a mini-cellulose format that enables storage of the encapsulated myoblasts and subsequent thawing to produce mini- myotube templates with viable cells that can be differentiated into elongated, striated, and multi- nucleated myotubes. Myotube templates derived from cryopreserved myoblast tissues exhibited comparable nuclei fusion indices, SAA coverage, and calcium transients to never frozen controls and enabled muscle stem cell engraftment to conduct a MEndR muscle injury and regeneration assay. We also performed a proof-of-concept demonstration that cryopreserved myoblast tissues could be shipped to domestic and international collaborators for subsequent use. Our novel cryopreservation strategy reduces the barrier for adoption of the mini-MEndR culture platform by circumventing the need for users to optimize the technically challenging steps of muscle template manufacturing, thus providing an easy-to-use, off-the-shelf platform for studying skeletal muscle health. More broadly, our cryopreservation approach offers a potential strategy to enable adoption of complex tissue engineered culture models beyond those for skeletal muscle.

## Acknowledgements

The authors would like to thank Kristyna Gorospe, Ting Yin, and Ruonan Cao for technical assistance. J.S and F.L.G acknowledge the CIQLE microscopy facility of SFR Santé Lyon-Est (UAR3453 CNRS, US7 Inserm, UCBL). Biorender was used to produce schematic elements.

## Author Contributions

S.R., E.J., B.X., S.K., J.S., H.L., N.G., and N.T.L. designed and performed experiments. S.R. and J.N. analyzed data and prepared figures. P.M.G., A.P.M., and S.R. conceived of the project. P.M.G., A.P.M., F.L.G., and B.Z. supervised the research. P.M.G., A.P.M., S.R., J.N., and H.L. wrote the manuscript. All authors reviewed and approved the manuscript.

## Data and Materials Availability Statement

The datasets generated during and analyzed in this study are available from the corresponding author on reasonable request.

