## Supplementary information for "A cryopreservation strategy for myoblast storage in paper-based scaffolds for inter-laboratory studies of skeletal muscle health"

This PDF file includes:

- Figures S1-S12
- Tables S1-S2
- Videos S1-S2
- References

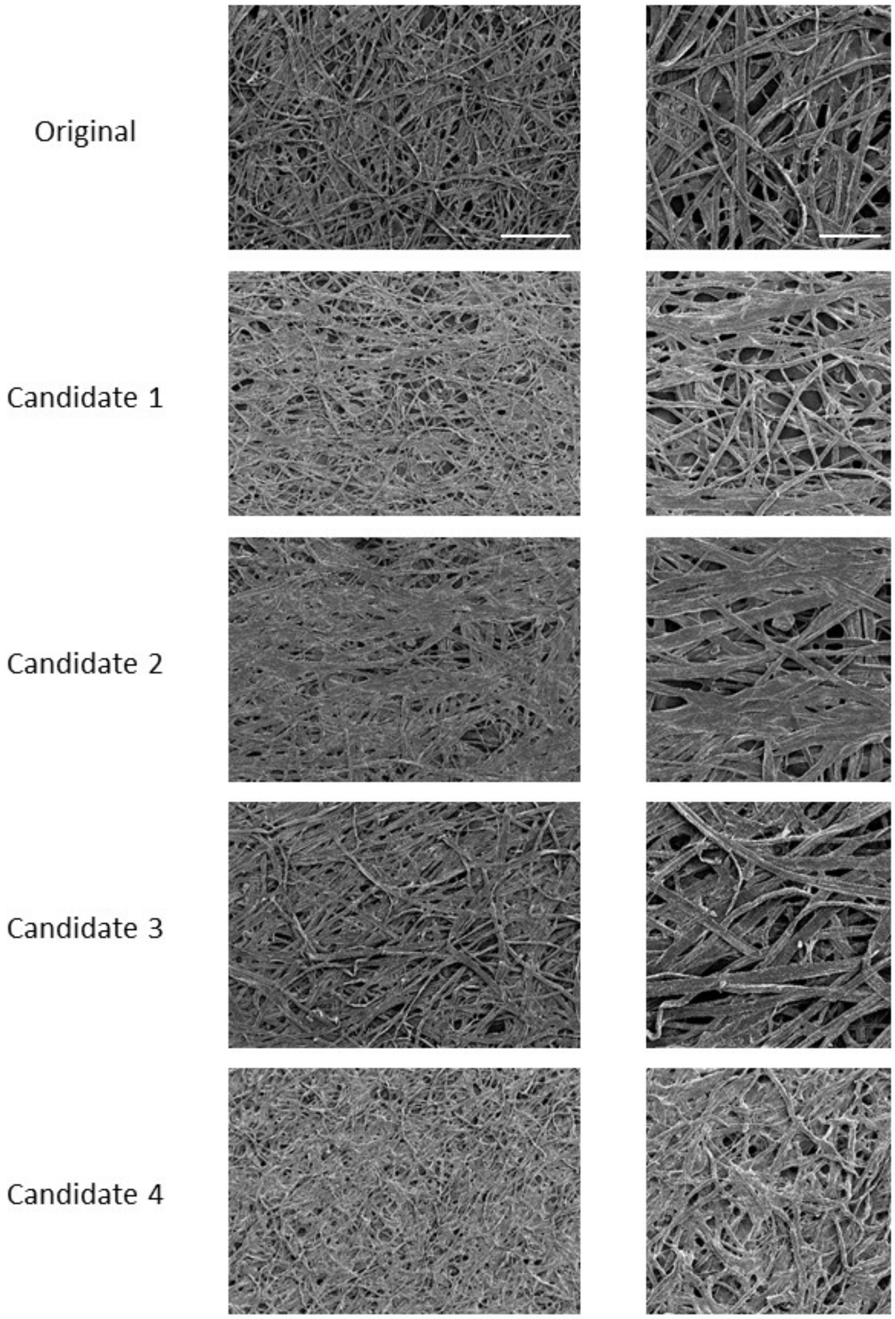


**Figure S1.** Representative 50X scanning electron microscopy (SEM) images of paper scaffold candidates. Cropped 800μm x 800μm sections taken from the center of the SEM images. Original = Mini-Minit, Candidate 1 = Finum, Candidate 2 = Teeli, Candidate 3 = Twin Rivers, Candidate 4 = Ahlstrom-Munksjo (see Methods). Note that the SEM image for Candidate 1 was used in Figure 1 and repeated here for comparison. Full image scale bar = 500μm; cropped image scale bar = 200μm.

A

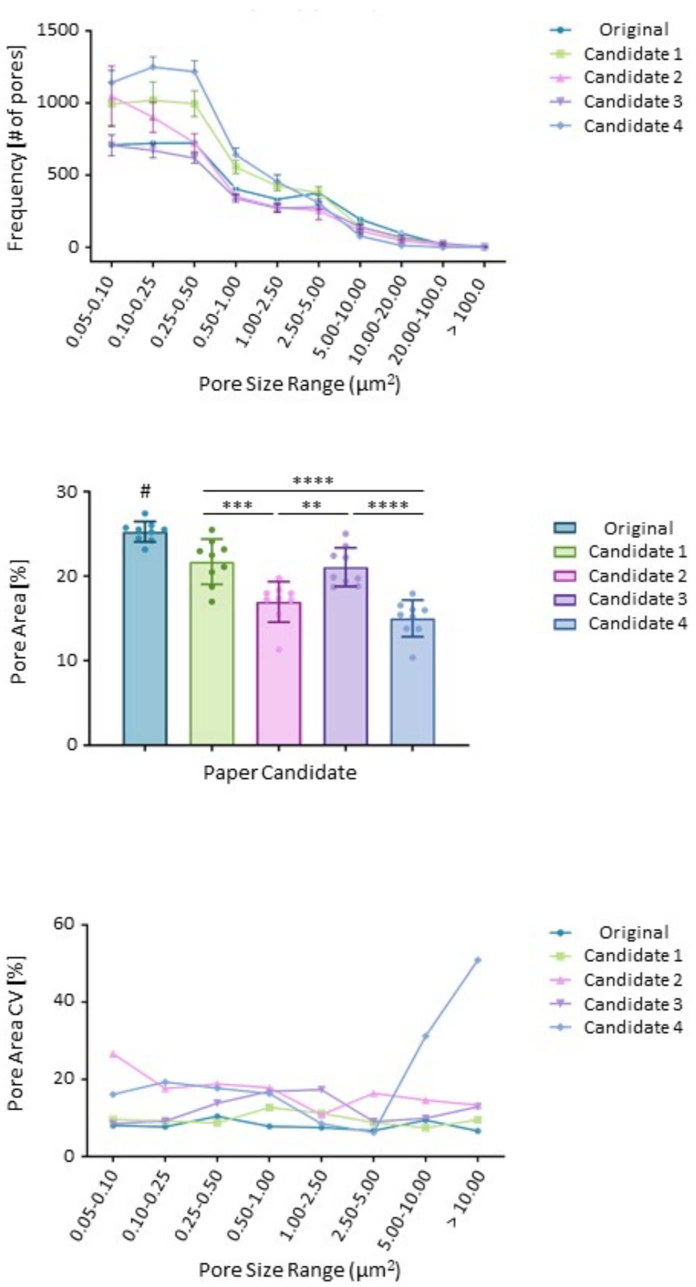


**Figure S2.** Porosity characterization of paper scaffold candidates. SEM images (Figure S1) were manually thresholded to distinguish between the pores and paper fibers. A list of all pores and their areas was generated using FIJI's Analyze Particles and aggregated as **(A)** histogram of the number of pores within a certain size range, **(B)** porosity (% area of the image occupied by the pores), and **(C)** histogram of the coefficient of variation (CV) of the area occupied by pores of a certain size.

Statistics performed using one-way ANOVA with Tukey post-test, #p<0.05 relative to all (*, ****, **, **** vs Candidates 1, 2, 3 and 4, respectively). *p<0.05 **p<0.01 ***p<0.001 ****p<0.0001. N=3 papers per candidate, n=3 sites per paper (9 total).

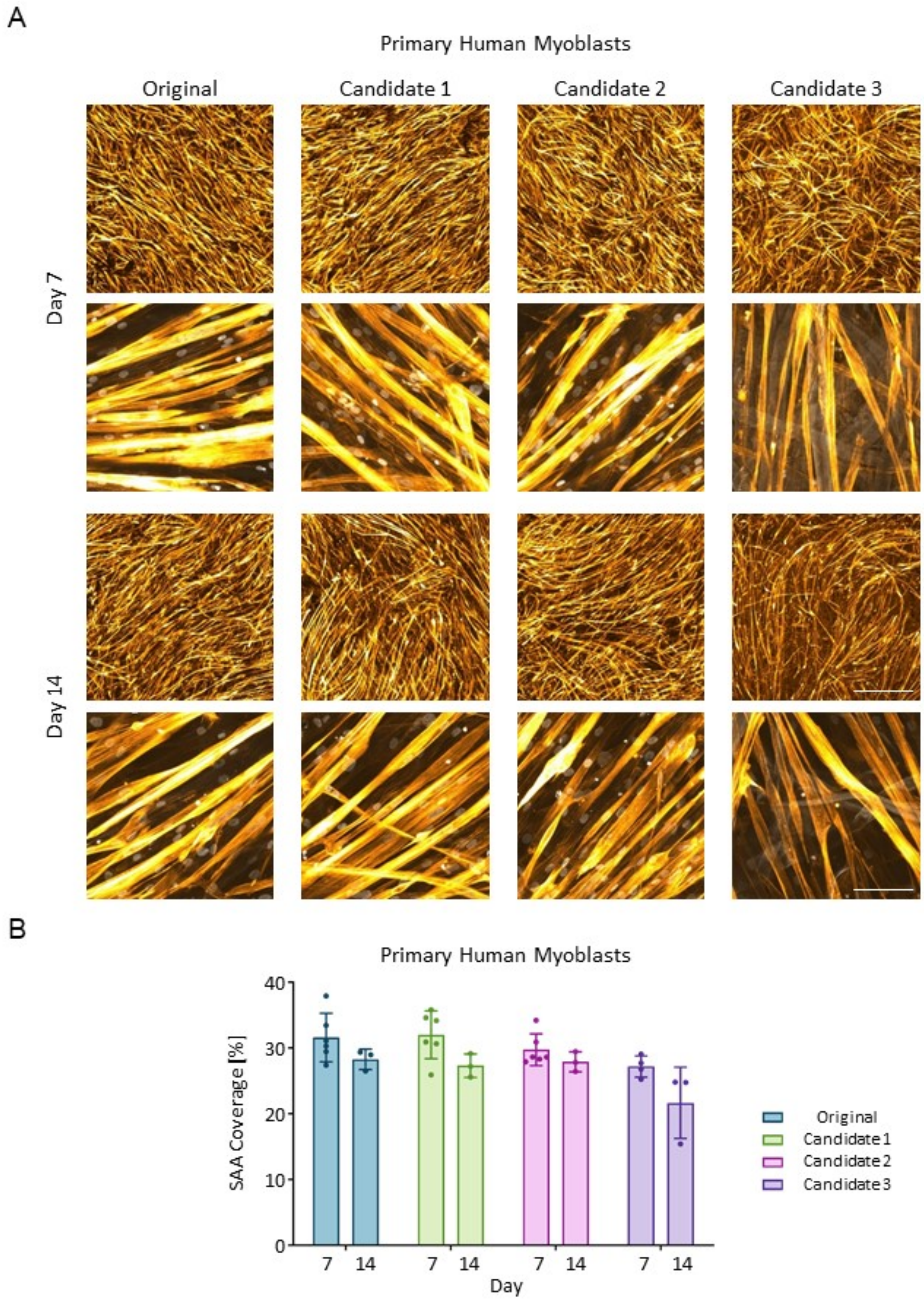


**Figure S3.** Analysis of primary human myoblast differentiation to myotubes in candidate cellulose scaffolds. Functional evaluation of paper scaffold candidates for differentiation of primary human myoblasts in cellulose scaffolds. **(A)** Representative 4X and 40X confocal images of myotube templates after 7 and 14 days of differentiation. Actin is shown in orange (Phalloidin), and nuclei are shown in grey (DRAQ5). 4X scale bar = 1000µm; 40X scale bar = 100µm. Note that the confocal images for primary human myoblasts in Candidate 1 were used in Figure 1 and repeated here for comparison. **(B)** Quantification of fiber area coverage of primary human myoblasts after 7 or 14 days of differentiation. Statistics performed using two-way ANOVA with Tukey post-test, ns – no significance. N=1-2 experimental replicates, n=3-6 technical replicates (scaffolds).

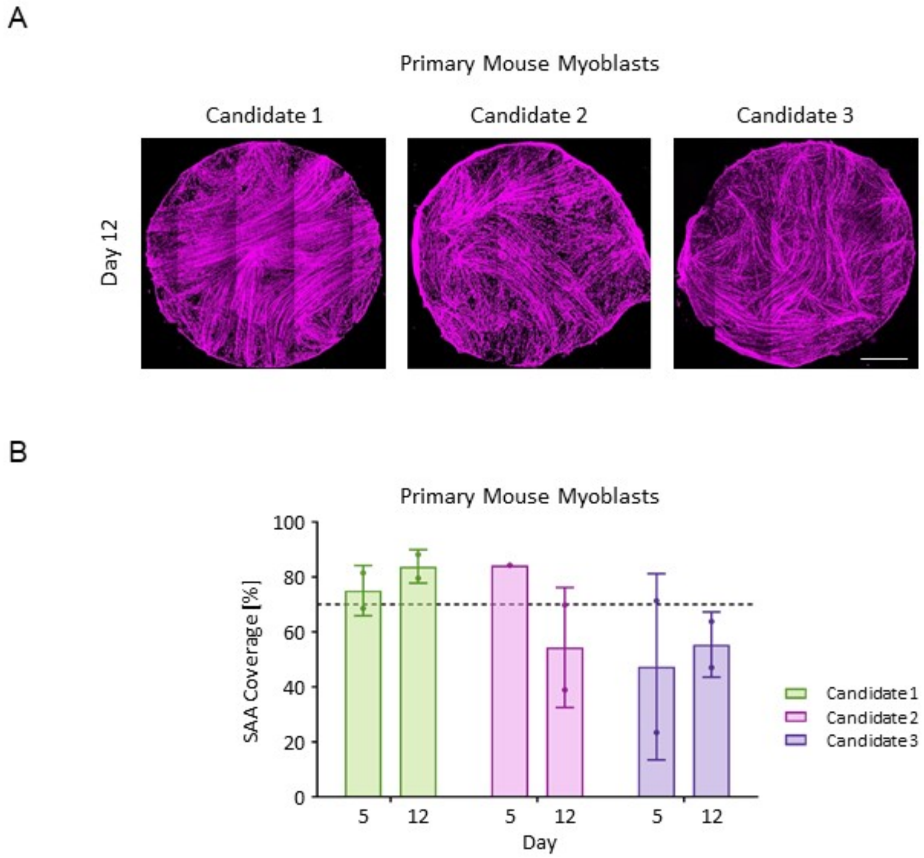


**Figure S4.** Analysis of primary mouse myoblast differentiation to myotubes in candidate cellulose scaffolds. **(A)** Representative 10X stitched confocal images of myotube templates after 12 days of differentiation. Sarcomeric-alpha-actinin (SAA) is shown in magenta. Scale bar = 1000µm. **(B)** Quantification of fiber area coverage of primary mouse myoblasts after 5 or 12 days of differentiation. Horizontal line represents the mean SAA coverage of primary mouse myoblasts seeded in the original paper candidate. Statistics performed using two-way ANOVA with Tukey post-test, ns – no significance. N=1 experimental replicate, n=1-2 technical replicates (scaffolds).

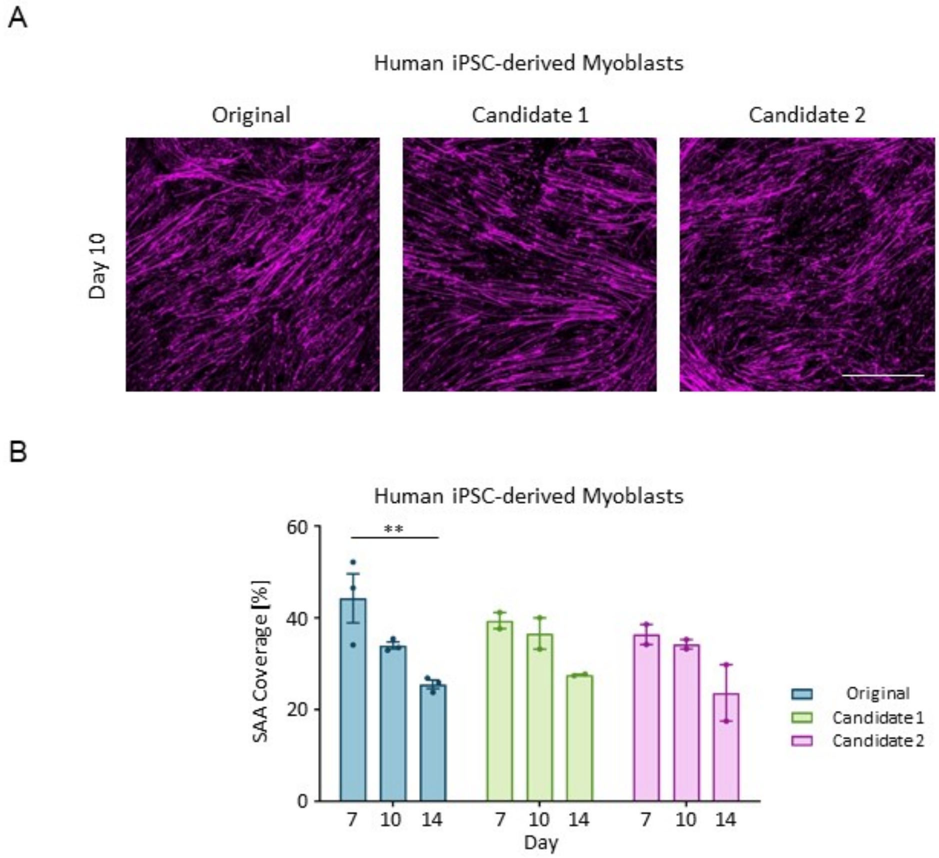


**Figure S5.** Analysis of human iPSC-derived myoblast differentiation to myotubes in candidate cellulose scaffolds. **(A)** Representative 4X confocal images of mini-MEndR myotube templates after 10 days of differentiation. Scale bar = 1000µm. Sarcomeric-alpha-actinin is shown (SAA) in magenta. **(B)** Quantification of fiber area coverage of human iPSC-derived myoblasts after 7, 10, and 14 days of differentiation. Statistics performed using two-way ANOVA with Tukey post-test, $^{**}p<0.01$. N=1-2 experimental replicates, n=2-3 technical replicates (scaffolds).

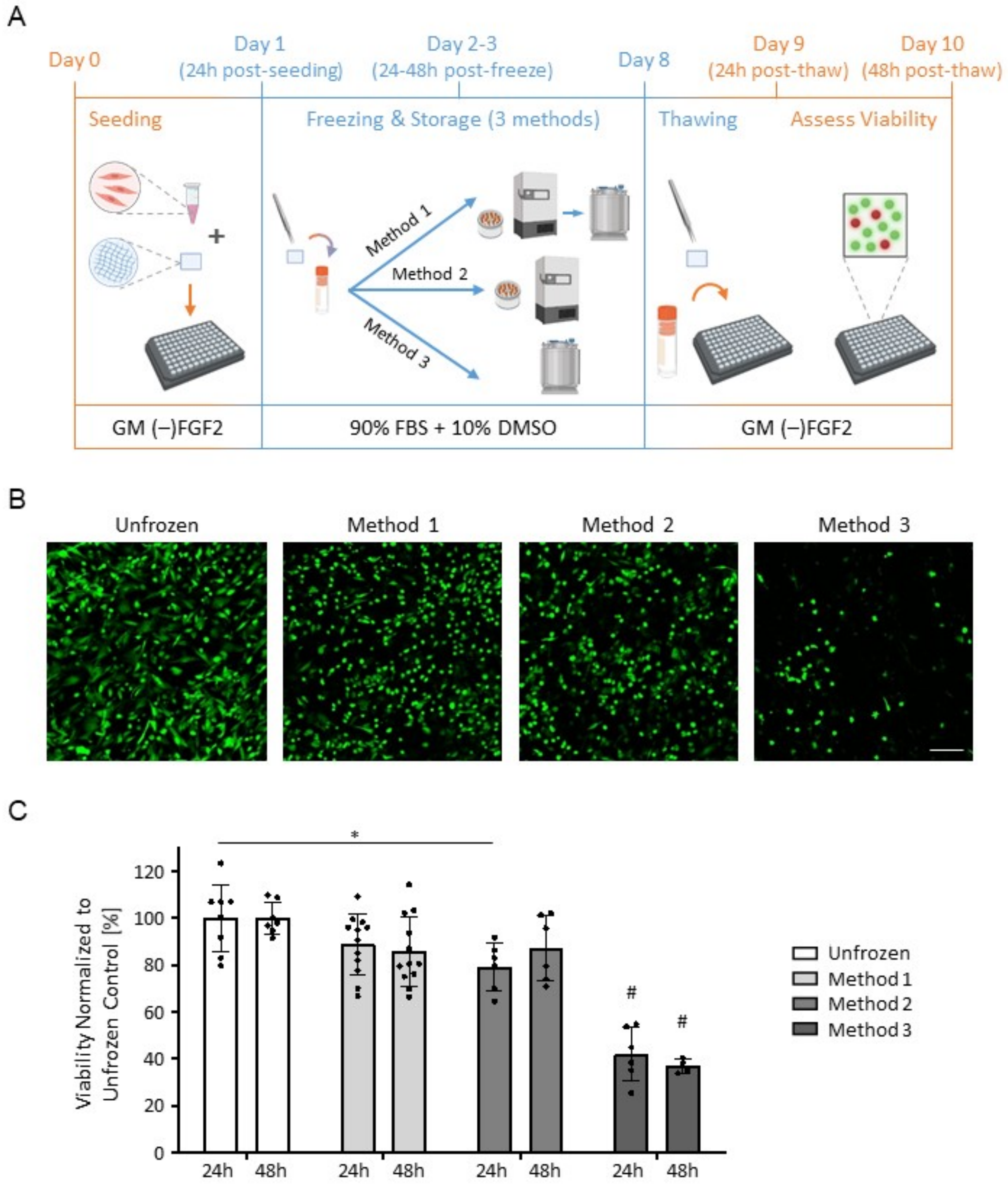


**Figure S6.** Evaluation of candidate protocols to freeze and store cellulose scaffolds seeded with hydrogel encapsulated myoblasts. **(A)** Schematic of tested protocols: Method 1: slow cooling in Nalgene® Mr. Frosty in -80°C freezer for 24-48h, followed by storage in liquid nitrogen for 5-6 days; Method 2: slow cooling in Nalgene® Mr. Frosty in -80°C freezer and maintained storage at -80°C for 1 week; or Method 3: direct cryovial storage in liquid nitrogen for 1 week. **(B)** Representative 20X live/total confocal images of primary human myoblasts in cellulose scaffolds either unfrozen, or 24h or 48h post-thaw with the mentioned freezing and storage methods. Live cells are shown in green (Calcein) and cell nuclei are show in grey (DRAQ5). **(C)** Quantification of cell viability (live cells / total cells) normalized to mean of unfrozen controls at the same timepoint. Cells were counted using a custom developed algorithm in FIJI (Figures S7-S9). Statistics performed using two-way ANOVA with Tukey post-test, #$p<0.0001$ relative to all conditions at the same timepoint, *$p<0.05$. N=1 experimental replicate, n=2 technical replicates (scaffolds), with s=2-4 sites each.

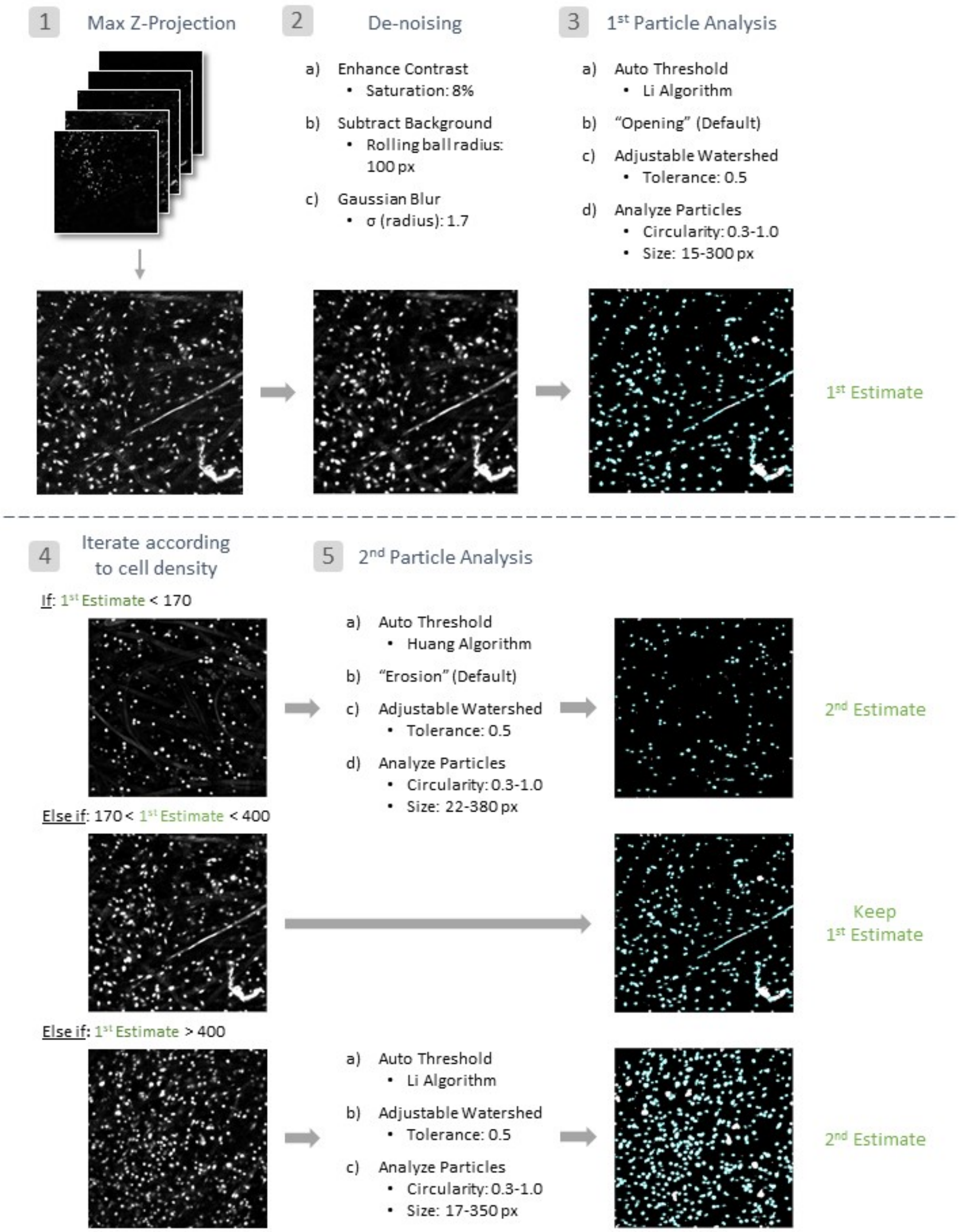


**Figure S7.** ImageJ algorithm for quantification of total cells. Input to the pipeline is a 20X image stack with a channel for nuclear staining (DRAQ5). Algorithm works as follows: (1) Stack of images captured at distinct depths projected at maximum intensity onto a single image plane (i.e. "Maximum Intensity Projection"); (2) background noise reduction using Subtract Background and Gaussian Blur, after optimization of pixel value saturation using Enhance Contrast function; (3)

1$^{st}$ particle analysis using Li algorithm for thresholding followed by an "Opening" and Adjustable Watershed step. The resulting mask is fed to the Analyze Particles function for automated cell counting; (4) classification of input image into low, medium or high cell density based on 1$^{st}$ estimate in step 3; (5) 2$^{nd}$ particle analysis based on results of (4), producing a final estimate of the number of total cells.

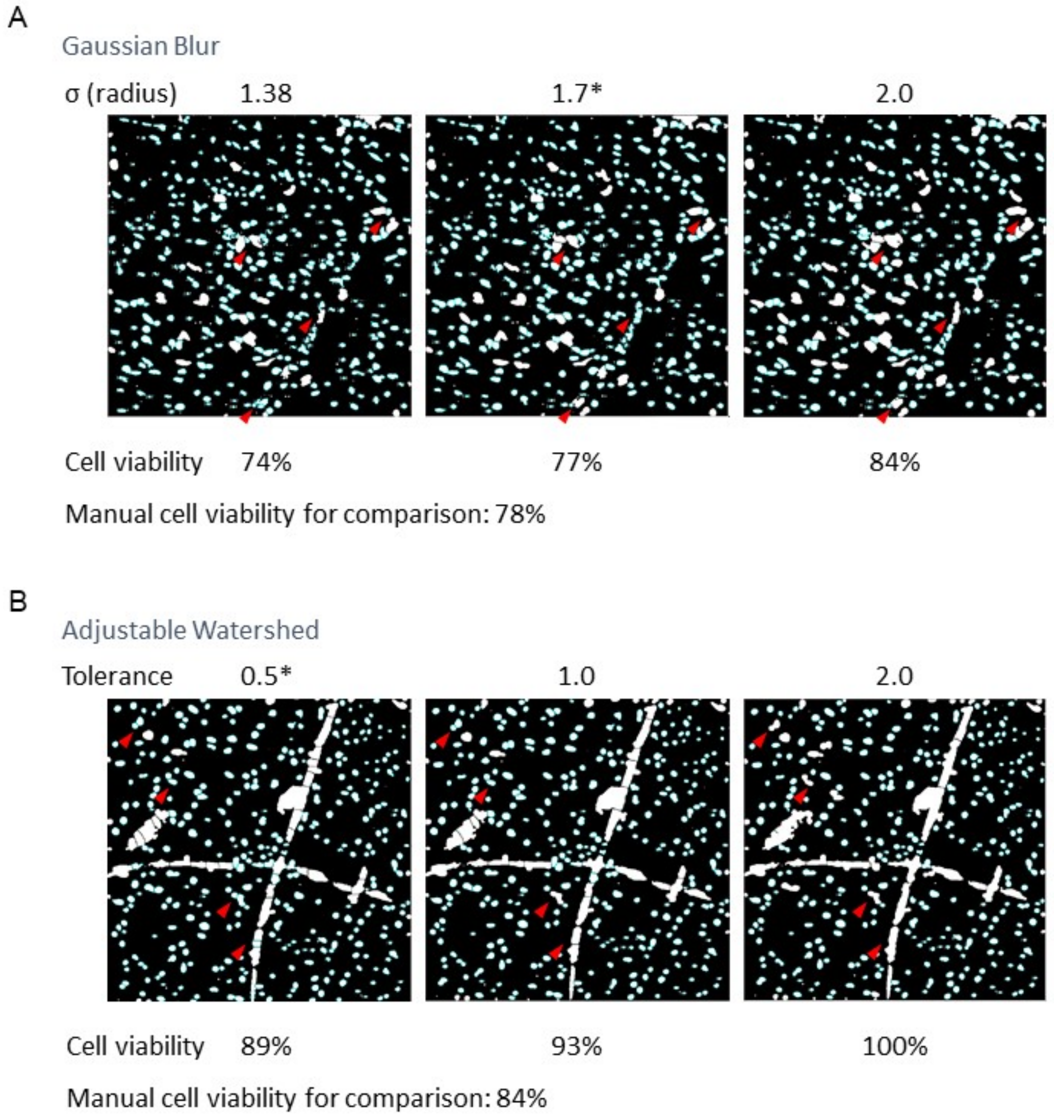


**Figure S8.** Optimization of Gaussian Blur radius and Adjustable Watershed tolerance for image analysis workflow to quantify live and total cells. The effect of tuning the parameters on cell viability was compared to manual annotation to select the optimal values. **(A)** Example of three Gaussian Blur sigma (radii) values (1.38, 1.7, 2.0), that were evaluated in their de-noising abilities. A value of 1.7 was determined to be optimal. **(B)** Example of three Adjustable Watershed tolerances (0.5, 1.0, 2.0), that were evaluated in their cell segmentation abilities. A value of 0.5 was determined to be optimal. Red arrows indicate regions of interest where subtle differences incurred by altering the parameters were demonstrated.

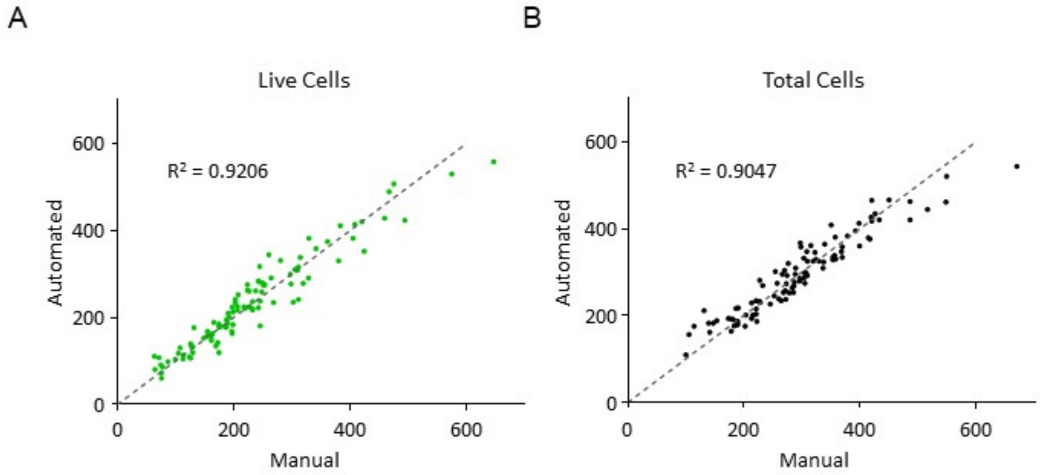


**Figure S9.** Accuracy of image analysis workflow by comparison to manual annotation. **(A)** Scatter plot of the number of live cells (Calcein channel) calculated by the automated algorithm in comparison to manual counting. The mean difference was 24 cells or 10.4%. R-squared=0.9206. **(B)** Scatter plot of the number of total cells (DRAQ5 channel) calculated by the automated algorithm in comparison to manual counting. The mean difference was 24 cells or 8.10%. R-squared=0.9047. Algorithm was tested with images encompassing a range of cell densities and observed to be relatively consistent. n=95 images quantified.

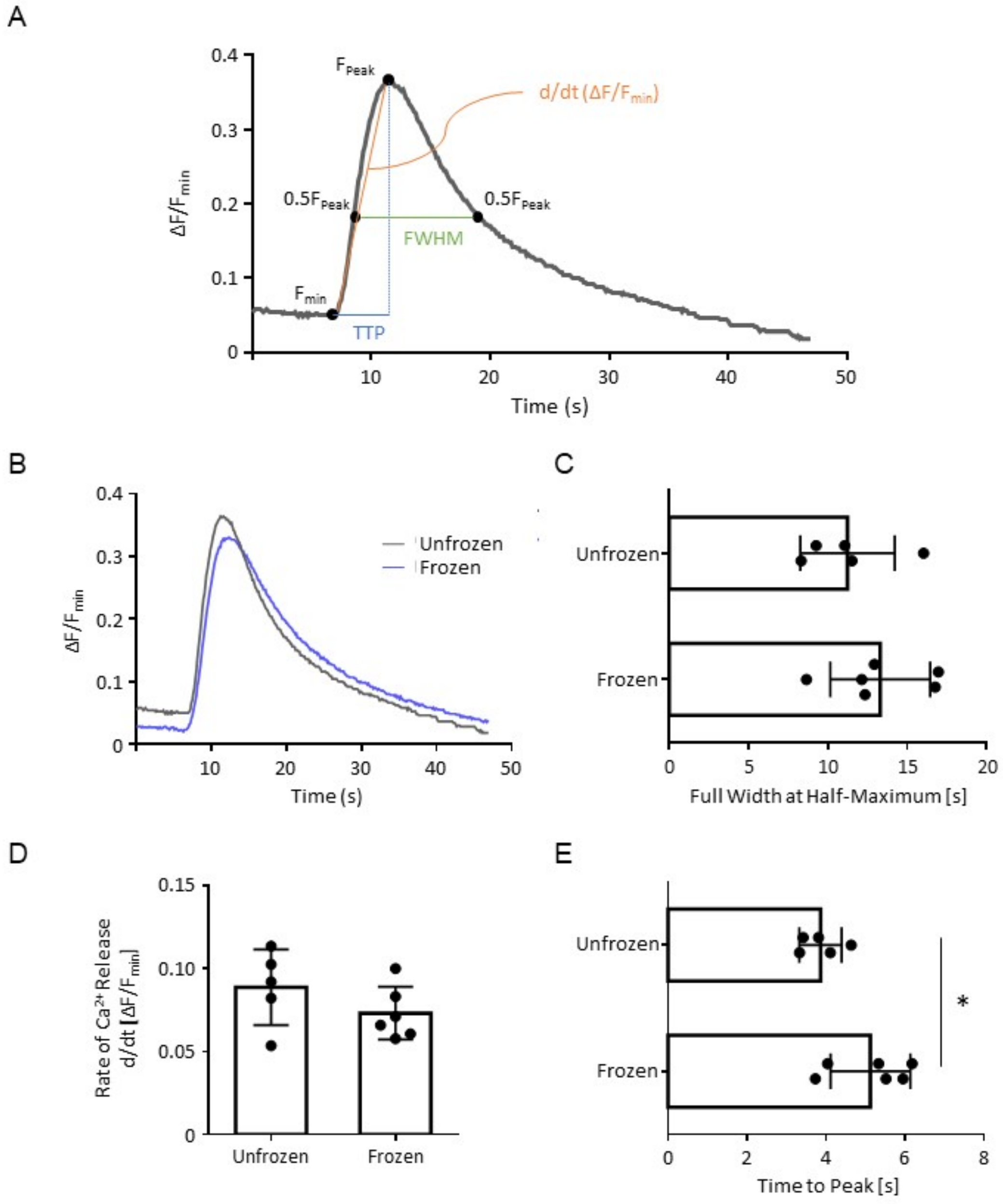


**Figure S10.** Evaluation of differentiation day 8 myotube calcium transients generated from primary human myoblasts previously cryopreserved in cellulose scaffolds. **(A)** Explanatory schematic of calcium handling metrics. First, relative fluctuations in the fluorescence were calculated and presented as $\Delta F/F0 = (F_{immediate} - F_{baseline})/(F_{baseline})$. Time to peak (TTP) is the time elapsed between the last datapoint of baseline fluorescence and the first datapoint at peak fluorescence. The slope of this line, $d/dt(\Delta F/F0)$, represents the rate of calcium release. The full-width at half-maximum (FWHM) represents the time elapsed between the normalized fluorescence first reaching half the peak fluorescence while ascending towards the peak until dropping below this value while descending from peak fluorescence. **(B)** Calcium transients of tissues after 8 days of differentiation. Curve represents average of N=1 experimental replicate, n=5-6 technical replicates (scaffolds). Image was presented in Figure 3 and repeated here to complement other metrics in this figure. **(C)** Rate of calcium release (quantified by the slope d/dt(ΔF/F0) of the ascending normalized fluorescence curve), **(D)** TTP, and **(E)** FWHM of calcium transients between unfrozen (never frozen) and frozen tissues. Statistics performed using unpaired two-tailed t-test, ns – no significance, *p<0.05. N=1 experimental replicate, n=5-6 technical replicates (scaffolds).

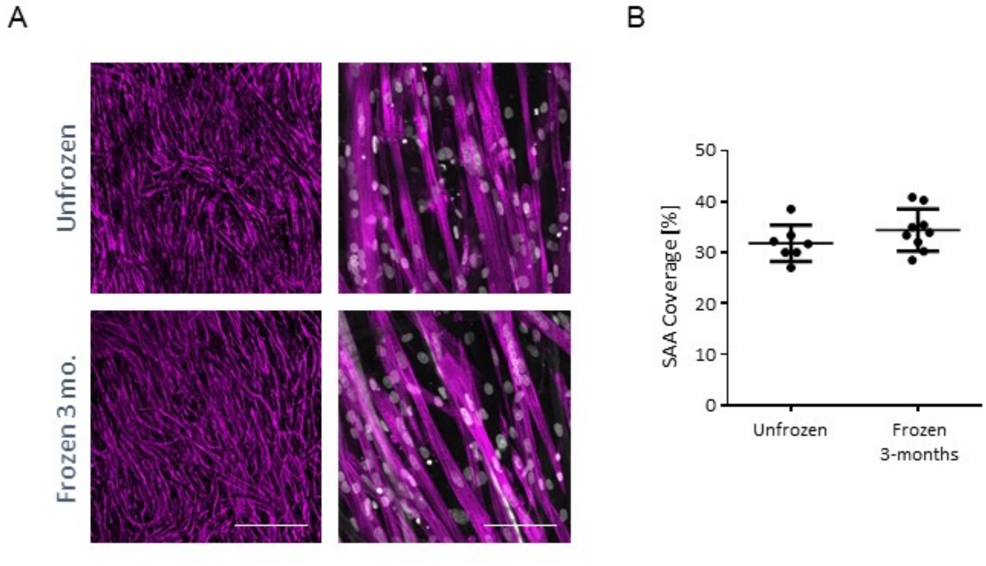


**Figure S11.** Cellulose scaffolds are a suitable vehicle for the long-term storage of cryopreserved myoblasts. **(A)** Representative 4X and 40X confocal images of primary human myoblasts differentiated for 7 days after being seeded in cellulose scaffolds and cryopreserved for 3 months in liquid nitrogen to investigate shelf life. Sarcomeric-alpha-actinin (SAA) is shown in magenta, and nuclei are shown in grey (DRAQ5). 4X scale bar = 1000µm; 40X scale bar = 100µm. **(B)** Bar graph quantification of SAA coverage from 4X images. Data also aggregated into Figure 3A and re-plotted here to visualize myotube template tissues derived from myoblasts stored for 3 months. Statistics performed using unpaired two-tailed t-test, ns – no significance. N=2 experimental replicate2, n=7-9 technical replicates (scaffolds).

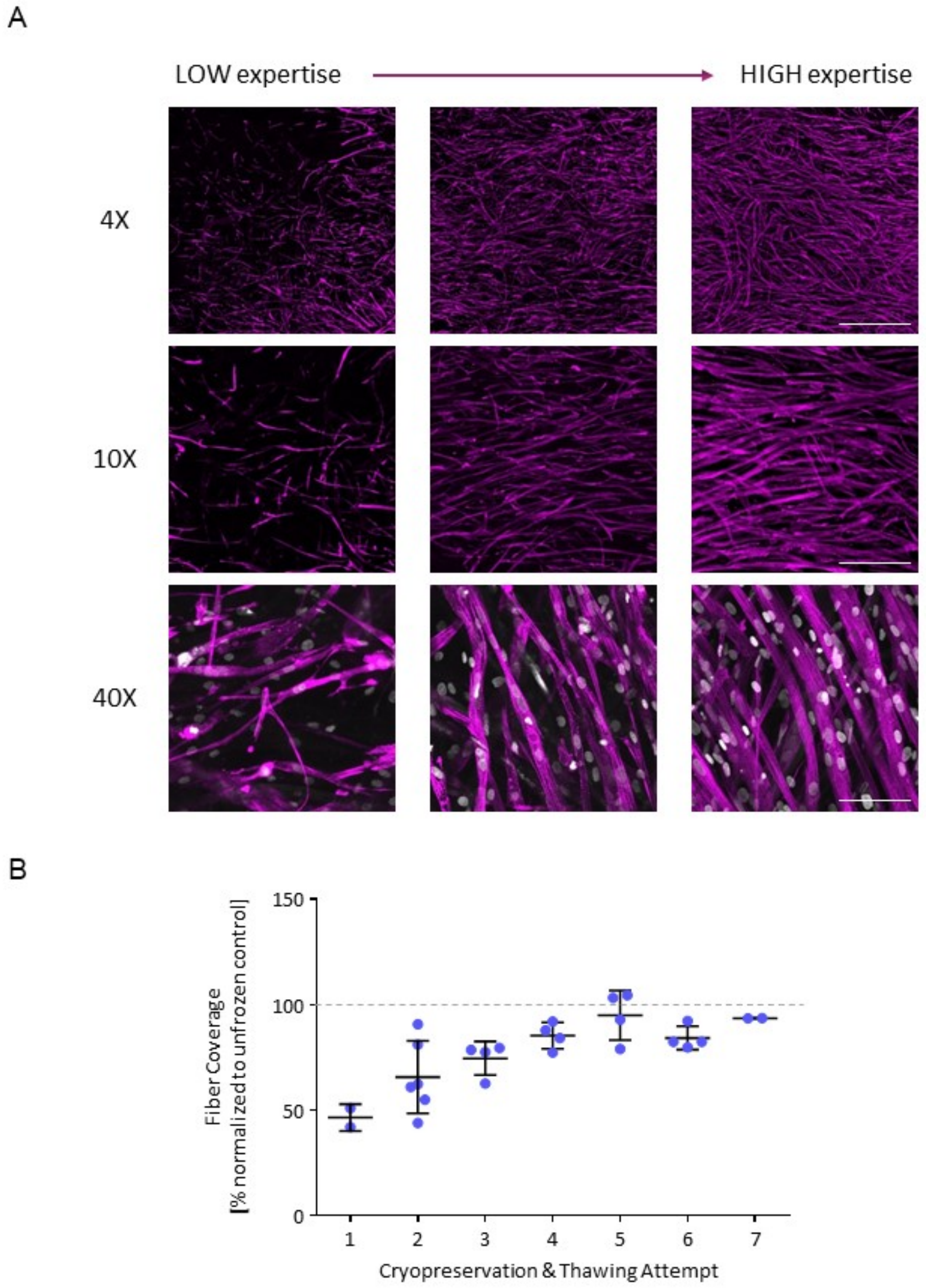


**Figure S12.** Cryopreservation SOP and subsequent myotube template cell culture can be quickly mastered by new users. **(A)** Representative confocal images of myotube templates generated by the same user at different levels of experience. Tissues were fixed after 7 days of differentiation and stained with SAA (magenta). Different magnification levels illustrate the effects on SAA coverage and myotube morphology. 4X scale bar = 1000µm; 10X scale bar = 400µm; 40X scale bar = 100µm. **(B)** Quantification of SAA tissue coverage attained by one user who trained from first-time user to expert level as indicated by plateauing coverage and homogeneous myotube templates. Coverage was normalized to the average SAA coverage from the unfrozen controls of each experiment.

**Table S1.** Parameters involved in the cryopreservation protocol, their current values, potential strategies for optimization, and priority level.

| Parameter | Current | Commentary & Potential for Optimization |
|---|---|---|
| Paper scaffold (Low priority) | Finum | Thin, cellulose-based, biocompatible paper scaffold. Already optimized as most comparable to original MEndR paper scaffold material for myotube template generation. However, comparison was in an unfrozen context only, so could compare templates derived from myoblasts frozen in different paper candidates. |
| Scaffold shape (Low priority) | Rectangle (0.5cm x 0.4cm) | Rectangular-shaped scaffolds are easily handled from their corners and demonstrated better alignment as myotubes were observed to align uniaxially to rectangle edges. Different rectangle dimensions that maintain an equal surface area can also be explored (e.g. 1.0cm x 0.2cm). |
| Seeding density (High priority) | 100k/tissue | Standard cell seeding density for generating primary human mini-myotube templates. Increased seeding density may help offset losses in cell recovery, although will reduce the number of tissues that can be fabricated. |
| Differentiation state (Low priority) | Undifferentiated | Only undifferentiated, but not differentiated myoblasts, attained expression of mature myogenic markers (e.g. MHC-1, MYF6) and myotube alignment by assay endpoint.[1] Nonetheless, the experiment could be explored in the mini-myotube template platform. |
| Freezing timepoint (Medium priority) | 24h post-seeding | Half-way through the 48h SM equilibration period in mini-myotube template protocol. Must balance stress from cryopreserving cells too soon after seeding (albeit assay and cell-type dependent [2-4]) and experiment duration (and tissue degradation over long-term). |
| Freezing method (Low priority) | Slow cooling 24-48h at -80°C in Nalgene® | Optimized (see Figure S6). Slow cooling also requires less CPA (toxicity risk), has a reduced risk of contamination, is less technically challenging, and is |

| | | |
|---|---|---|
| | Mr.Frosty, then liquid nitrogen | amenable to larger sample sizes such as mini-myotube templates.[5] |
| Freezing medium / Cryoprotective agent (High priority) | 90% FBS + 10% DMSO | Both are field standards.[6-7] Future experiments could also explore combinations of CPAs with DMSO (see Section 2.5.3, e.g. trehalose, glycols, etc.). Freezing medium temperature can also be measured to elucidate the extent of its influence (given that DMSO has a temperature-dependent cytotoxicity). |
| Freezing volume (Low priority) | 1 mL | Larger volumes should offer more buffer to temperature perturbations, but will take longer to thaw (increasing exposure to CPA toxicity and thermal gradients). Pilots are suggested (e.g. 0.5 mL). |
| Thawing method (Low priority) | Rapid thawing in 37°C water bath | Standard in the field.[8] Pilots demonstrated that thawing too many cryovials in water bath at once and being unable to remove tissues from CPA quickly enough led to significant cell death (Figure S12). However, limitation is throughput. Also, to avoid re-opening the liquid N2 tank for each cryovial, a subset of cryovials are taken at a time and kept cool in a frozen Nalgene® Mr. Frosty in an ice bucket. Monitoring the temperature within the Nalgene® Mr. Frosty can provide data into the efficacy of this set-up. |
| Thawing medium (Medium priority) | Seeding Medium (SM) | Same as prior to freezing. All commercially available for easy adoption by collaborators. However, could explore media additives that have been shown to improve cell viability and other hallmarks when added to post-thawing media, such as ROCKi,[9] cysteine cathepsin inhibitors,[1] or anti-oxidants.[10] |
| Equilibration time (Medium priority) | 24h | Half-way through the 48h SM equilibration period in myotube template protocol, although not optimized rigorously in this study. Must balance stress on cells induced from thawing and experiment duration (and tissue degradation over long-term). |

**Table S2.** Breakdown of experimental replicates and statistical analyses.

| Figure | Number of experimental replicates (N) | Number of technical replicates tissues/wells (n) | Number of images per tissue/well | Statistical Test |
|---|---|---|---|---|
| 2C | 4 | Unfrozen: 8<br>Frozen: 9 | 3-4 | Unpaired two-tailed t-test |
| 3C | 5 | Unfrozen: 17<br>Frozen: 23 | 1 | Unpaired two-tailed t-test |
| 3D | 3 | Unfrozen: 11<br>Frozen: 22 | 2<br>(plotted avg.) | Unpaired two-tailed t-test |
| 4B | 2 | Unfrozen: 8<br>Frozen: 10 | 1-2 | Unpaired two-tailed t-test |
| 5D | 1 | In-house: 6<br>Domestic: 5<br>International: 2 | In-house: 2<br>Domestic: 1-3<br>International: 3<br>(plotted avg.) | Ordinary one-way ANOVA on tissue averages with Tukey's multiple comparison's test |
| 5E | 1 | In-house: 6<br>Domestic: 4<br>International: 2 | In-house: 2<br>Domestic: 2<br>International: 4<br>(plotted avg.) | Ordinary one-way ANOVA on tissue averages with Tukey's multiple comparison's test |
| 6C | 3 | Unfrozen:<br>DMSO CTX+: 11<br>P38i CTX+: 12<br>Frozen:<br>DMSO CTX+: 9<br>P38i CTX+: 11 | 21 stitched images/tiles | Unpaired two-tailed t-test p38i vs control for each of unfrozen and frozen |
| S2B | 1 | All: 3 | 3 | Ordinary one-way ANOVA with Tukey's multiple comparison's test |
| S3B | Day 7: 2<br>Day 14: 1 | Day 7:<br>Candidate 3: 4<br>All others: 6<br>Day 14:<br>All: 3 | 1 | Two-way ANOVA with Tukey's multiple comparison's test |
| S4B | 1 | Candidate 1: 2<br>Candidate 2: 1-2<br>Candidate 3: 2 | 1 | Two-way ANOVA with Tukey's multiple comparison's test |
| S5B | 1 | Original: 3<br>Candidate 1: 2<br>Candidate 2: 2 | 1 | Two-way ANOVA with Tukey's multiple comparison's test |

| | | | | |
|---|---|---|---|---|
| S6C | 1 | Unfrozen: 2<br>Method 1: 2<br>Method 2: 2<br>Method 3: 2 | Unfrozen: 4<br>Method 1: 3<br>Method 2: 3<br>Method 3: 2 | Two-way ANOVA with Tukey's multiple comparison's test |
| S10C | 1 | Unfrozen: 5<br>Frozen: 6 | 1 | Unpaired two-tailed t-test |
| S10D | 1 | Unfrozen: 5<br>Frozen: 6 | 1 | Unpaired two-tailed t-test |
| S10E | 1 | Unfrozen: 5<br>Frozen: 6 | 1 | Unpaired two-tailed t-test |
| S11B | 2 | Unfrozen: 7<br>Frozen: 9 | 1 | Unpaired two-tailed t-test |

**Supplementary Videos**

https://u.pcloud.link/publink/show?code=kZ21kF0ZByFocj0JQYpy9YYYJaU9fjCwNxBV

**Video S1.** Representative 4X fluorescence intensity video of calcium transients from baseline, to peak fluorescence, and return to baseline, for control myotube template. Primary human myoblasts differentiated over 8 days were incubated with Calbryte$^{TM}$ 520 calcium indicator dye and stimulated with 2mM Acetylcholine. Scale bar = 200µm. See Figure S10 for explanation of video-based characterizations.

**Video S2.** Representative 4X fluorescence intensity video of calcium transients from baseline, to peak fluorescence, and return to baseline, for frozen myotube template. Primary human myoblasts were cryopreserved in cellulose scaffold, then thawed and differentiated over 8 days and incubated with Calbryte$^{TM}$ 520 calcium indicator dye and stimulated with 2mM Acetylcholine. Scale bar = 200µm. See Figure S10 for explanation of video-based characterizations.
